# RNA m^6^A demethylase ALKBH4 governs whitefly development

**DOI:** 10.64898/2026.08.08.743705

**Authors:** Jing Yang, Chao Wang, Peipan Gong, Chao He, Buli Fu, Shaonan Liu, Xuegao Wei, Cheng Yin, Mingjiao Huang, Tianhua Du, Jinjin Liang, Xuguo Zhou, Ralf Nauen, Youjun Zhang, Chris Bass, Xin Yang

## Abstract

N6-methyladenosine (m^6^A) modification is the most predominant and ubiquitous internal modification of RNA in eukaryotes, serving as a key post-transcriptional regulator of gene expression that is dynamically modulated by methyltransferases (writers) and demethylases (erasers). However, while the functions of m^6^A methylases have been partially elucidated in insects, the identity of m^6^A erasers in arthropods and their chemical catalytic mechanisms, as well as biological functions, remains largely enigmatic. Here, we uncovered 2499 putative methylase genes and 1148 putative demethylase genes in 266 insect genomes, and demonstrated that ALKBH4 functions as an m^6^A demethylase in the whitefly, *Bemisia tabaci*, catalyzing the oxidative reversal of mRNA m^6^A modifications both in vitro and in vivo. Furthermore, we established that ALKBH4, in coordination with other core components of the m^6^A pathway, fulfills an essential function in regulating the transcript stability of Imaginal Disk Growth Factor 1 (IDGF1) during whitefly development. Collectively, our findings expand the evolutionary scope of the eukaryotic m^6^A modification system, and reveal a conserved yet insect-specific epitranscriptomic regulatory mechanism governing fundamental physiological processes and adaptive phenotypes.

**Significance statement:** The addition of a methyl group to the N6-position of adenosine (m^6^A) is a highly abundant chemical modification of RNA. However, the functional role of m^6^A in insects and the key enzymes that regulate its levels remains poorly understood. In this study, we explored putative methylase genes and demethylase genes in hundreds insect genomes, and identified an m^6^A RNA demethylase, ALKBH4, in the whitefly, *Bemisia tabaci*. We demonstrate that ALKBH4 oxidatively reverses mRNA methylation in vivo and in vitro, in combination with other components of the m^6^A pathway, plays an important role in whitefly development. These findings provide new insight into m^6^A methylation system of insect.

## Introduction

N6-methyladenosine (m^6^A) modification is the most predominant modification of mRNA, and plays crucial roles in modulating diverse biological processes through its action on gene expression, mRNA stability, translation, splicing, and pri-miRNA processing. m^6^A methylation predominantly occurs at the conserved DRACH motif (D: G/A/U/C; R: G/A; H: U/A/C), which is enriched near stop codons, the 5’ and 3’ untranslated regions (UTRs), and within long internal exons in mammals (1–8). In mammalian systems, m^6^A modification is a reversible process, that requires methylases (writers), demethylases (erasers), and binding proteins (readers). The m^6^A writer complex, which catalyzes m^6^A methylation, is composed of two core enzymes: methyltransferase-like 3 (METTL3) that contributes the catalytic residues (9) and methyltransferase-like 14 (METTL14) that stabilizes the structure of the complex through methyltransferase domains (10). These core subunits act in combination with several auxiliary cofactors, including WT1 associated protein (WTAP, 11), KIAA1429 (12), RNA binding motif protein 15 (RBM15, 13), methyltransferase-like 16 (METTL16, 14) and CCCH type 13 zinc finger protein (ZC3H13, 15) to modulate methylation activity. Importantly, m^6^A methylation can be erased by m^6^A demethylases, all of which are members of the alpha ketoglutarate-dependent dioxygenase (ALKB) family of Fe(II)/α-ketoglutarate(α-KG)-dependent dioxygenases. Fat-mass and obesity-associated protein (FTO) was the first m^6^A demethylase to be identified in mammals (16), and plays important roles in development, neurogenesis and tumorigenesis (17–20). ALKBH5 is another mammalian demethylase reported to impact RNA metabolism and mouse fertility (21). Finally, ALKBH10B is an m^6^A demethylase in *Arabidopsis*, affecting floral transition and vegetative growth (22).

In insects, m^6^A writers and readers have been linked to the regulation of a range of biological functions. In *Drosophila*, comprehensive molecular and physiological characterization of the components of the methyltransferase complex has revealed their important roles in neuronal functions, sex determination, and pre-mRNA splicing (23–25). In *Bombyx mori*, METTL3, METTL14, and YTHDF3 may regulate the response to nuclear polyhedrosis virus by m^6^A modification (26). In *Tribolium castaneum*, m^6^A methylases regulate ecdysis during eclosion and reproduction (27). In *B. tabaci*, methyltransferases have been implicated in the regulation of the P450 gene *CYP4C64*, increasing its expression and resulting in resistance to thiamethoxam (28). Recently, a candidate m^6^A eraser, ALKBH8, was identified in *Aedes aegypti* and *Drosophila melanogaster*, and incubation of recombinant ALKBH8 with total RNA was found to result in a reduction in global m^6^A levels in dot blot assays using an anti-m^6^A antibody (29). Similarly, ALKBH5 was identified as a potential m^6^A demethylase in *Locusta migratoria*, where it was linked to the regulation of aggregation behavior (30). However, our understanding of m^6^A demethylases in most insect species and their biological role(s) remains limited. Importantly, chemical verification of the activity of insect m^6^A demethylases *in vitro* has not been demonstrated.

The whitefly, *B. tabaci* (Hemiptera: Aleyrodidae), is a globally distributed, highly damaging pest of agriculture that attacks a wide range of food and commodity crops (31). *B. tabaci* is a species complex of at least 30 cryptic species, some of which (e.g., Mediterranean [MED] and Middle East-Asia Minor 1 [MEAM1]) are devastating crop pests (32). The intensive use of insecticides to manage this insect pest has led to the evolution of widespread resistance (33–35). As noted above, resistance to the insecticide thiamethoxam mediated by *CYP4C64* in this species has been linked to the action of methyltransferases on m^6^A binding sites (28). However, the role of m^6^A demethylases in other biological processes in *B. tabaci*, remains unknown. Here, we identify ALKBH4, as an m^6^A demethylase in *B. tabaci* and demonstrate its capacity to reverse m^6^A modifications *in vivo* and *in vitro*. We demonstrate the role of this demethylase and the m^6^A pathway more generally in regulating nymph development in *B. tabaci*. Our results advance our understanding of the reversible nature of m^6^A modifications and their functional roles in insects.

## Results

### m^6^A “writers” and “erasers” in 266 insect species

To systematically identify putative m^6^A “writers” and “erasers” across the diversity of insects, 266 insect genomes were interrogated for the presence of genes encoding putative m^6^A methylases and demethylases based on the presence of conserved domains present in these genes. This resulted in the identification of 2,499 putative methylase genes and 1,148 putative demethylase genes. The mean number of putative methylase genes identified in the insect genomes analysed was nine, ranging from six in the silkworm, *Bombyx mori* to 20 in the click beetle, *Ignelater luminosus*. The mean number of putative demethylase genes identified was four, ranging from one in *Anopheles atroparvus* to eight in the monarch butterfly, *Danaus plexippus* (Fig. 1A, S2A, Table S1, Dataset S1, S2).

**Fig. 1.**
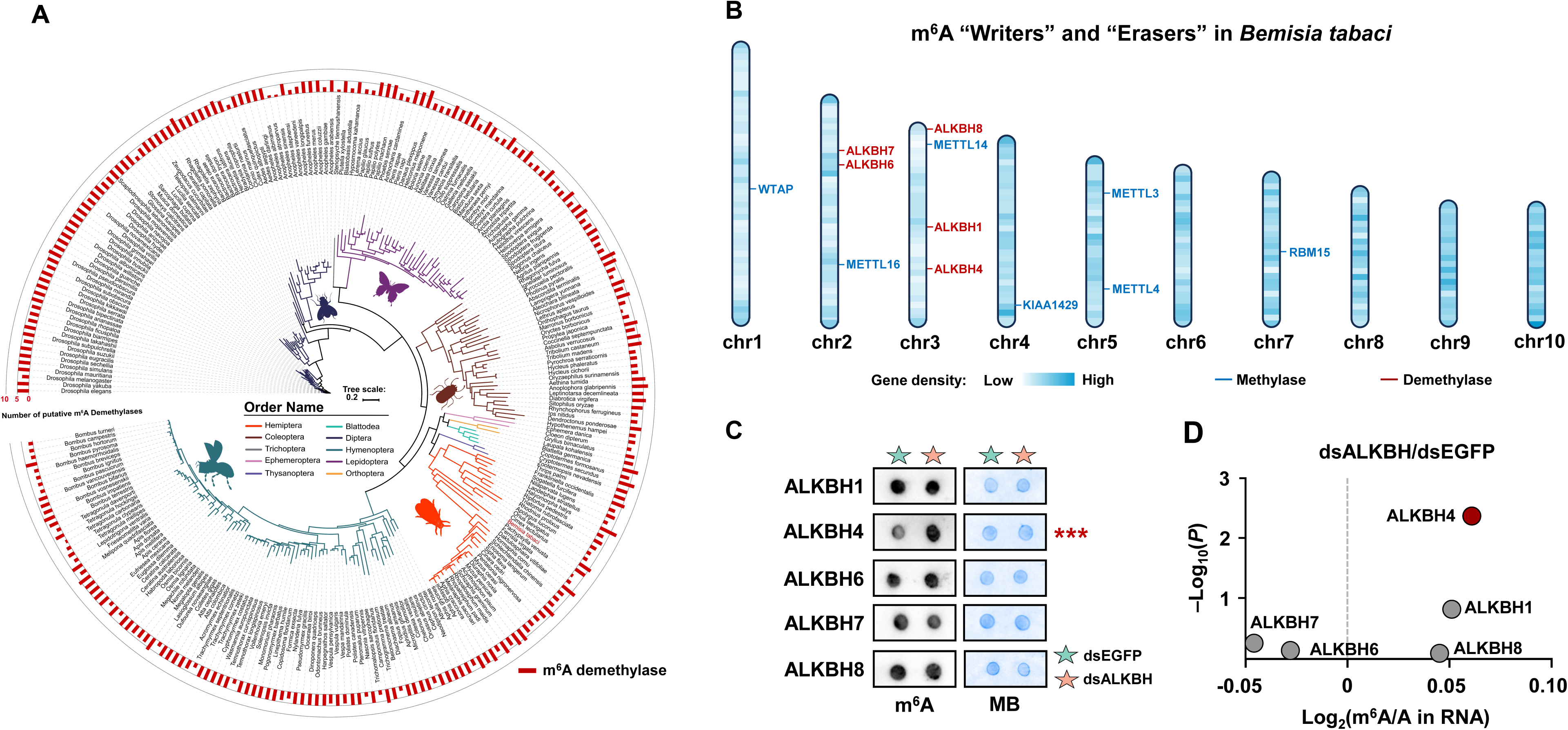
Characterization of insect m^6^A methylase and demethylase genes. (A) Number of putative m^6^A methylase and demethylase genes in 266 insect species. Gene numbers are presented on a maximum likelihood phylogeny of the species. Methylase genes are homologous genes of *METTL3*, *METTL14* (PF05063), *METTL16* (PF05971), *WTAP* (PF17098), *KIAA1429* (PF15912), *RBM15* (PF00076); Demethylase genes are homologous genes of *ALKBH4* (PF03171), *ALKBH5* (PF13532) and *FTO* (PF12933). (B) Chromosomal location of whitefly m^6^A methylase and demethylase genes in *B. tabaci*. (C) Dot blot assays of global m^6^A levels in RNA extracted from *B. tabaci* fed dsRNA corresponding to different putative m^6^A demethylase genes or dsEGFP. MB: methyl blue. (D) The m^6^A/A ratio of total RNA, determined by UPLC‒MS/MS, derived from *B. tabaci* fed dsRNA corresponding to different putative m^6^A demethylase genes.

### The m^6^A methylation pathway of *B. tabaci*

Among the m^6^A methylase and demethylase genes identified above, seven methylase and five demethylase genes were both found in the whitefly, *B. tabaci* MED and MEAM1. As shown in Fig. 1B, the m^6^A “writers” METTL3/4, WTAP, KIAA1429, METTL14, METTL16 and RBM15 are located on chromosome 5, 1, 4, 3, 2 and 7, respectively. The m^6^A “erasers” ALKBH1/4/8, and ALKBH6/7 are located on chromosome 3 and 2, respectively. Intriguingly, all five of these m^6^A demethylase genes were found to be intronless (Fig. S1, Table S2).

To investigate the involvement of methylases in whiteflies, ultra-performance liquid chromatography-tandem mass spectrometry (UPLC-MS/MS), a technique that combines high-performance liquid chromatography for separating compounds with mass spectrometry for quantifying specific molecules, was used to quantify m^6^A/A levels following RNA interference (RNAi)-mediated knockdown (KD) of each of seven candidate methylase genes. Relative mRNA levels of seven genes were significantly downregulated after feeding with dsRNAs (*METTL3*: decreased by 43%, *P =* 9.62 × 10^-5^; *METTL14*, 50%, *P =* 4.18 × 10^-5^; *WTAP*, 40%, *P =* 0.0038; *KIAA1429*: 54%, *P =* 0.0002; *RBM15*, 44%, *P =* 0.0046; *METTL4*, 52%, *P =* 0.0013; *METTL16*, 41%, *P =* 0.0014, Fig. S2B). UPLC-MS/MS analysis revealed a significant increase in m^6^A levels following interference with each whitefly methylase, compared to the EGFP control group (*METTL3*: *P =* 0.0004; *METTL14*, *P =* 4.17 × 10^-5^; *WTAP*, *P =* 0.0022; *KIAA1429*: *P =* 0.0016; *RBM15*, *P =* 0.0110; *METTL4*, *P =* 0.0005; *METTL16*, *P =* 0.0032, Fig. S2C).

To examine whether the five m^6^A demethylase genes are involved in whitefly m^6^A modification, dot blot assays, a method for detecting specific RNA modifications on membranes, and UPLC-MS/MS were used to detect global m^6^A levels after RNAi knockdown of each of the demethylase candidate genes. Feeding *B. tabaci* adults with dsRNA corresponding to each demethylase gene resulted in a significant decrease in mRNA expression levels of all five genes (*ALKBH1*: decreased by 39%, *P =* 1.69 × 10^-4^; *ALKBH4*: 43%, *P =* 4.49 × 10^-5^; *ALKBH6*: 50%, *P =* 5.70 × 10^-9^; *ALKBH7*: 40%, *P =* 9.10 × 10^-5^; *ALKBH8*: 32%, *P =* 9.19 × 10^-4^, Fig. S3A). Dot blot analysis of total m^6^A levels in RNA derived from each of the five RNAi treatments using a specific m^6^A antibody revealed that only silencing of *ALBKH4* was associated with a significant increase in m^6^A levels (Fig. 1C, S3B). This finding was validated by UPLC-MS/MS quantification of m^6^A/A ratios in total RNA of each treatment/control, which showed significant increases in m^6^A/A ratios in *ALKBH4* knock-down treatments compared to the dsEGFP control (*P =* 0.008, Fig. 1D). Together, these results provide evidence that whitefly *ALKBH4* acts as an m^6^A demethylase *in vivo*.

### ALKBH4 demethylates m^6^A-modified RNA *in vitro*

To further investigate the role of ALKBH4 as an mRNA demethylase (Fig. 2A), we recombinantly expressed this protein in *E*. *coli* and performed demethylation activity assays with the recombinant protein. A synthetic 15-mer m^6^A-modified single strand RNA (ssRNA, 0.5 nmol, AUUGUCA**m^6^A**CAGCAGC, 16, Table S7) was used as a substrate in assays which contained an equal amount of the recombinant full-length ALKBH4 protein (0.5 nmol). ALKBH4 was found to demethylate the m^6^A-containing ssRNA with increasing activity over the first 10 minutes with the reaction reaching saturation in 30 minutes (Fig. 2B, S2C), and the demethylation level reaching 50% (Fig. 2C). The ALKB family proteins require iron ligand residues and α-KG ligand residues for their enzymatic activity. The mutant ligands demonstrated a lower binding affinity for the protein compared to the native ligand (Fig. S4). To further validate these findings and gain insight into domains of ALKBH4 that are key to its activity, we tested the demethylation activity of mutant versions of ALKBH4 where we modified two key domains (ALKBH4-MU1: H169A/ D171A where two iron (II) ligand residues were mutated and ALKBH4-MU2: R257Q/ R263Q where two α-KG ligand residues were mutated, Fig. 2D, E, S3C). These mutants exhibited a significant reduction in m^6^A-demethylation activity (ALKBH4-MU1:12.7-fold, *P =* 8.55 × 10^-6^; ALKBH4-MU2: 32.4-fold, *P =* 1.79 × 10^-7^) compared to the wild type (ALKBH4-WT) (Fig. 2F, G). Together, these results demonstrate that ALKBH4 catalyzes oxidative demethylation of m^6^A in an iron- and α-KG-dependent manner.

**Fig. 2.**
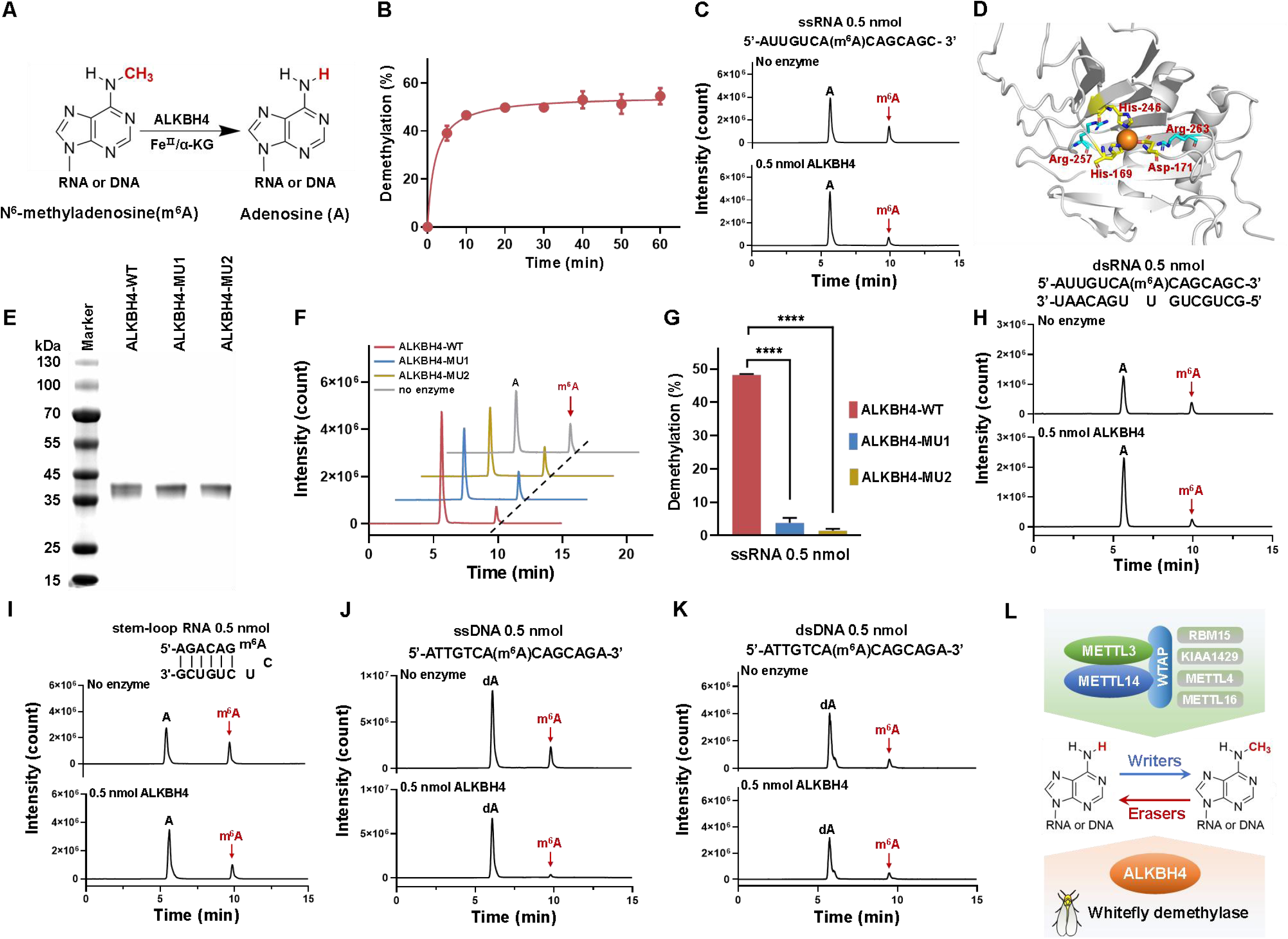
Oxidative demethylation of m^6^A in nucleic acids by ALKBH4. (A) Schematic of the proposed oxidative demethylation of m^6^A to adenosine in RNA or DNA by ALKBH4 in the presence of Fe(II) and α-KG. (B) ALKBH4-mediated demethylation (%) of an m^6^A-modified 15-mer ssRNA over time (*n* = 3, means ± SE). All reactions were performed at pH 7.0 and 25°C. (C) ALKBH4 (0.5 nmol) demethylation of m^6^A in ssRNA (0.5 nmol) at pH 7.0 and 25 °C for 30 min as revealed by UPLC‒MS/MS analysis of the digested substrates. (D) 3D structure of the *B. tabaci* ALKBH4 protein. (E) Heterologous expression of the ALKBH4 wild type (ALKBH4-WT) and mutant proteins (ALKBH4-MU1/2) in *E*. *coli* as detected by SDS-PAGE. (F) Demethylation of an m^6^A-modified 15-mer ssRNA at pH 7.0 and 25°C by recombinant ALKBH4-WT and ALKBH4-MU1/2 (0.5 nmol). (G) The demethylated ratio of ALKBH4-WT and ALKBH4-MU1/2 proteins (*n* = 3, mean ± SE, \*\*\*\**P* < 0.0001, two-tailed Studentʹs *t* test). (H, I, J, K) ALKBH4 (0.5 nmol) demethylated m^6^A in dsRNA (H), stem-loop RNA (I), ssDNA (J), dsDNA (K) at pH 7.0 and 25 °C for 30 min as reveled by UPLC‒MS/MS analysis of the digested substrates. (L) A schematic of the m^6^A pathway in *B. tabaci*. Seven m^6^A methylase genes and one demethylase gene are involved in the m^6^A pathway in this species.

To examine the demethylation activity of ALKBH4 towards a series of m^6^A-containing synthetic oligonucleotides, we performed mass spectrometry, under the same reaction conditions and observed demethylation yields of 36% for dsRNA (Fig. 2H), 35% for stem-loop RNA (Fig. 2I), 86% for ssDNA (Fig. 2J), and 32% for dsDNA (Fig. 2K). Together, these results provide unequivocal evidence that ALKBH4 confers demethylation activity toward m^6^A-modified ssRNA, dsRNA, stem-loop RNA, ssDNA and dsRNA. They also suggest that the m^6^A pathway in *B. tabaci* involves at most seven m^6^A methylase genes and one demethylase gene (Fig. 2L).

### Different binding sites effect the activity of m^6^A methylation and demethylation

To understand the nature and frequency of m^6^A binding sites in the genome of *B. tabaci* we interrogated the genome for 24 different permutations of the consensus m^6^A binding site DR<u>A</u>CH (D = C/A/U/G; R = A/G; H = A/C/U) and compared the results with the patterns of m^6^A sites observed in the genomes of *Homo sapiens* and *Arabidopsis thaliana* (Fig. 3A). This analysis identifies conserved m^6^A motifs in the genome and compares their distribution across species. A total of 20,748, (an average of 62.7%) of genes contained m^6^A binding sites in *B. tabaci*, ranging from 47.1% for the CGACC site to 80.2% for the AAACA site; a total of 27,655, (average of 66.4%) of genes in *A. thaliana*, ranging from 37.1% (CGACC) to 89.6% (AAACA); and total of 21,507, (average of 79.8%) of genes In *H. sapiens*, ranging from 56.0% (CGACU) to 86.1% (GGACA) (Table S3). Intriguingly, these m^6^A binding sites are most abundant in gene coding sequence (CDS) in *B. tabaci* and *A. thaliana*, whereas in *H. sapiens*, m^6^A sites are most abundant in the CDS and 3’ untranslated region (UTR) (Fig. 3B, Table S4, Dataset S3, S4, S5). To investigate the action of methylases and demethylases at these sites, METTL3, FTO, and ALKBH5 from *H. sapiens*, METTL3 and ALKBH10B from *A. thaliana*, and METTL3 and ALKBH4 from *B. tabaci* were functionally expressed *in vitro* and their activity against 24 ssRNAs representing permutations of the DRACH site (Table S3) assessed (Fig. S3D, E). The methylation activity status of the three METTL3 enzymes from different species were similar, with the last nucleic acid (A) of the motif found to play a key role in influencing activity, and the binding site CAACA most highly methylated (Fig. 3C, Table S5). The demethylation activity of human FTO was close to 100% for the majority of the ssRNA substrates. In contrast, the demethylation activity of ALKBH subfamily of *H. sapiens*, *A. thaliana* and *B. tabaci*, was less efficient, with the activity for most of ssRNA substrates lower than 50% (Fig. 3D, Table S6).

**Fig. 3.**
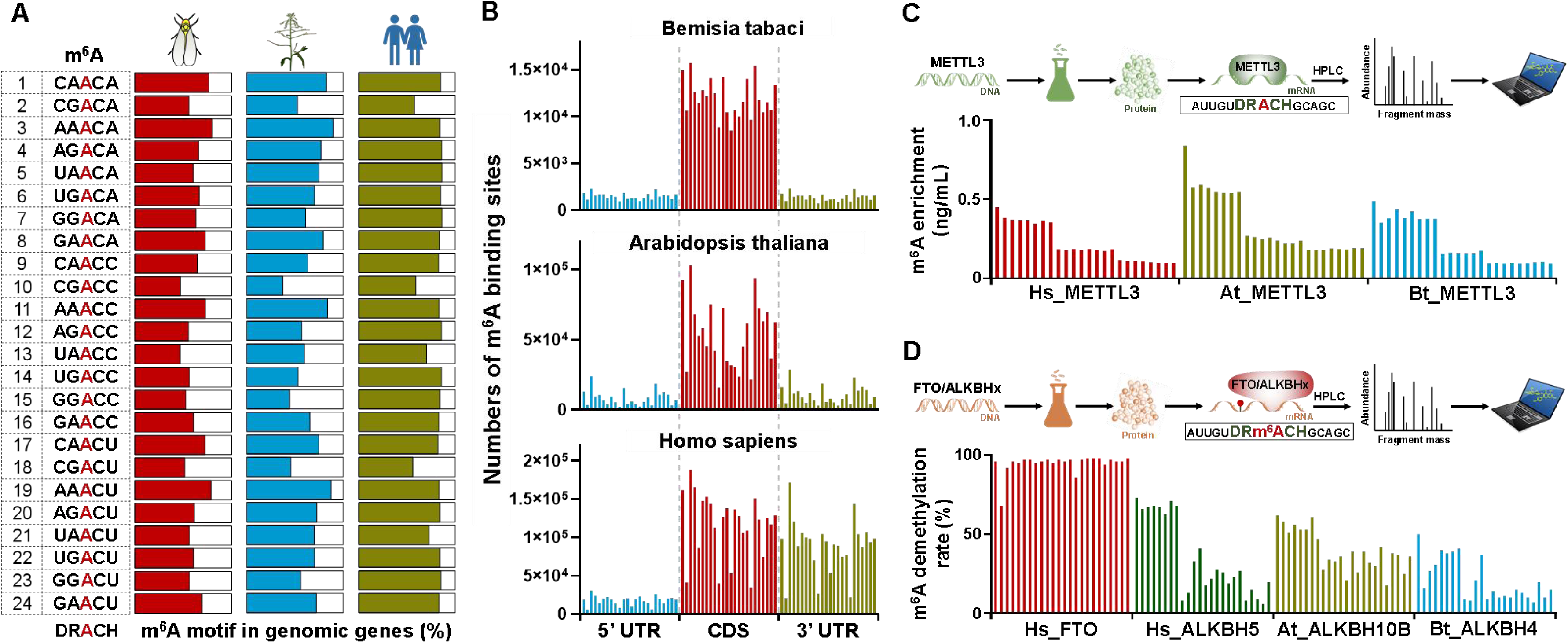
m^6^A binding site sequence influences methylation and demethylation activity. (A) The percentage of m^6^A consensus binding sites (24 forms of DRACH: D represents G/A/U/C; R represents G/A; H represents U/A/C) in genomes of *B. tabaci*, *A. thaliana*, and *H. sapiens*. (B) Number of m^6^A binding sites in the 5’ UTR, CDS or 3’ UTR of genes in *B. tabaci*, *A. thaliana*, and *H. sapiens*. (C) The methylation activity of METTL3 of three species (Hs_ METTL3: METTL3 of *H. sapiens*; At_ METTL3: METTL3 of *A. thaliana*; Bt_ METTL3: METTL3 of *B. tabaci*) on 24 forms of m^6^A substrate. (D) The demethylation activity of four demethylase genes (Hs_FTO: FTO of *H. sapiens*; Hs_ALKBH5: ALKBH5 of *H. sapiens*; At_ALKBH10B: ALKBH10B of *A. thaliana*; Bt_ALKBH4: ALKBH4 of *B. tabaci*) from three species on 24 forms of m^6^A substrate.

### m^6^A regulates *IDGF1* and its role in nymph development in *B. tabaci*

To identify candidate genes regulated by ALKBH4, transcriptome sequencing of *B. tabaci* was performed after RNAi knockdown of *ALKBH4*. RNA sequencing provides a high-throughput method to measure gene expression and identify differentially expressed genes after perturbation. Differential gene expression analysis identified 224 genes as significantly up-regulated upon silencing of *ALKBH4*, and 27 genes as down-regulated (Fig. 4A, Dataset S6), Gene ontology enrichment analysis revealed terms associated with tetrapyrrole binding, oxidoreductase activity and fatty acid metabolism as significantly enriched in genes upregulated upon silencing of *ALKBH4* (Fig. S5). Interestingly, transcriptome sequencing analysis revealed a differentially expressed gene, *Imaginal Disk Growth Factor 1* (*IDGF1*), which is strongly associated with insect development and biological regulation. IDGFs are chitinase-like secretory proteins that play a key role in insect development, and CRISPR-CAS knock-out of IDGF genes in *Drosophila melanogaster* significantly lowers viability and fertility (35).

**Fig. 4.**
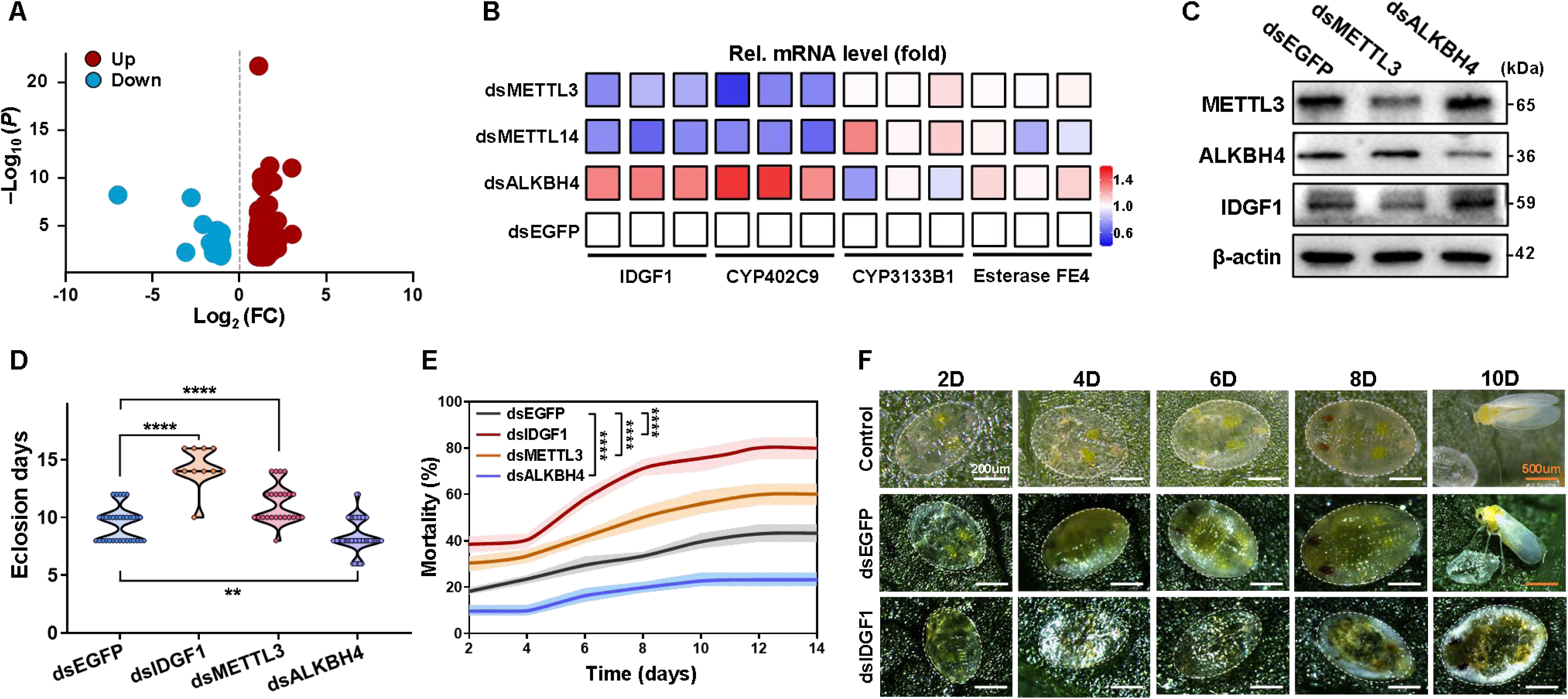
The m^6^A pathway modulates whitefly nymph development by methylating a chitinase mRNA. (A) Volcano plot showing differentially expressed genes identified in RNA-seq analysis following *ALKBH4* knockdown in *B. tabaci* (red, overexpression; blue, underexpression. (B) Relative mRNA level of genes involved in nymph development and insecticide resistance following knockdown of the m^6^A methylase genes *METTL3* and *METTL14* or the demethylase gene *ALKBH4*, as detected by qRT-PCR. (C) Western blot analysis of the expression levels of METTL3, ALKBH4, IDGF1 after RNAi knockdown *METTL3* and *ALKBH4*. β-Actin was used as a loading control. (D, E) Time to eclosion (days) and mortality of *B. tabaci* after RNAi knockdown of *IDGF1*, *METTL3* or *ALKBH4* every 48 h from the second nymphs until death (\*\**P* < 0.01, \*\*\*\**P* < 0.0001, two-tailed Studentʹs *t* test). (F) Representative images displaying development of *B. tabaci* nymphs after RNAi knockdown of *IDGF1*. Nymph stage: scale bars, 200 μm. Adult stage: scale bars, 500 μm.

To investigate the functional role of m^6^A modification in the regulation of candidate gene, *IDGF1,* we used quantitative real time polymerase chain reaction (qRT-PCR) to examine the effect of knockdown of the m^6^A methylase genes *METTL3* and *METTL14*, and the m^6^A demethylase gene *ALKBH4* on its expression. The expression of *IDGF1* was found to be significantly downregulated following knockdown of *METTL3* and *METTL14* (*IDGF1*: dsMETTL3, *P =* 0.0059; dsMETTL14, *P =* 0.0114), and significantly upregulated following knockdown of *ALKBH4* by qRT-PCR and western blot analyses (*IDGF1*: *P =* 0.0029, Fig. 4B, C).

Given the important physiological role of chitinase-like secretory family, and our finding that knockdown of *ALKBH4* results in the downregulation of *IDGF1*, we were motivated to examine the role of the m^6^A pathway in regulating this gene. The full-length cDNA sequence of *IDGF1* contained a 1542-bp ORF encoding 513 amino acid residues (Fig. S6). To explore the role of *IDGF1* in the development of *B. tabaci*, we examined the time taken for nymphs to develop to adults after RNA interference of *IDGF1*. The expression of *IDGF1* was significantly reduced by 60% after feeding second instar nymphs with dsRNA corresponding to this gene for 48 h (*P =* 1.33 × 10^-5^, Fig. S7A). After RNAi knockdown of *IDGF1*, the days taken to reach eclosion and mortality were significantly increased (eclosion days: *P =* 8.67 × 10^-12^, mortality: *P =* 7.36 × 10^-13^, Fig. 4D, E, and F). Moreover, the developmental time of the second, third and fourth nymphal instar stages were also significantly increased (second stage, *P =* 5.08 × 10^-6^; third stage, *P =* 0.0071; fourth stage, *P =* 0.0346, Fig. S7C). To further validate the developmental role of IDGF1, we used a plant-mediated virus-induced gene silencing (VIGS) system to continuously suppress IDGF1 throughout development. Continuous silencing of IDGF1 significantly delayed egg hatching (*P* = 2.64 × 10⁻⁴, Fig. S8A, B), prolonged nymphal development (*P* = 2.49 × 10⁻⁴, Fig. S8C), and increased cumulative mortality (*P* = 6.62 × 10⁻⁴, Fig. S8D). Consistent with the RNAi results, VIGS confirmed that IDGF1 is essential for normal development of *B. tabaci*, and that its silencing causes developmental delay and arrest (Fig. S8E).

To examine the functional role of METTL3 and ALKBH4 in *B. tabaci* development, the mortality and developmental duration of nymphal stages was monitored following RNAi knockdown of *METTL3* or *ALKBH4*. The relative mRNA expression levels of *METTL3* or *ALKBH4* were significantly decreased by 52% and 59%, respectively after feeding second instar nymphs with dsRNA corresponding to this gene for 48 h (*P =* 1.29 × 10^-5^; *P =* 0.0016, Fig. S7A). A significant increase in days taken to reach eclosion (*P =* 8.45 × 10^-6^) and mortality (*P =* 9.24 × 10^-6^) was observed following knockdown of *METTL3* expression. In contrast, a significant decrease in days taken to reach eclosion (*P =* 3.20 × 10^-3^) and mortality (*P =* 4.11 × 10^-9^) were exhibited following knockdown of *ALKBH4* (Fig. 4D, E).

To investigate if putative m^6^A binding sites are present in *IDGF1* the mature mRNA sequence of this gene was interrogated for the m^6^A consensus motif DRACH. The DRACH motif is a conserved sequence recognized by m^6^A methylases in RNA. Seven high-confidence sites were predicted in the *IDGF1* mRNA with two m^6^A binding sites in the 5’ untranslated region (UTR) and five sites in the coding sequence (CDS) (Fig. S9). According to the location of these sites, we constructed a sequence fragment containing two putative m^6^A binding sites located in the 5’ UTR (A-51, A-109) and cloned these into the pGL4.26 reporter gene plasmid (Fig. 5A). A second set of constructs (IDGF1 G-51&G-109) were created where the putative m^6^A binding sites in each of the two IDGF1 A-51&A-109 fragments was mutated (A mutated to G in all cases). These constructs were then transfected into *D. melanogaster* S2 cells and their expression levels examined in reporter gene assays. Reporter gene assays are used to evaluate the activity of a gene or promoter by measuring the expression of a reporter gene. No significant differences of reporter gene activity were found between IDGF1-MU1, IDGF1-MU2, IDGF1-MU3 and IDGF1-MU4, compared to the respective wild type controls (Fig. 5A). To confirm the role of m^6^A binding sites in the CDS in modulating gene expression, we generated constructs where the m^6^A consensus sequences observed in the CDS of *IDGF1* were changed from DR<u>A</u>CH to DR<u>G</u>CH and cloned into the pAC5.10 plasmid. Cell lines expressing the IDGF1-G1191 construct exhibited a significant (*P* = 1.63 × 10^-4^) reduction in gene transcription compared with the line expressing the wild-type IDGF1-A1191 construct (Fig. 5B), with western blot analysis confirming that the encoded protein is significantly decreased in the mutant construct as a result (Fig. S7D).

**Fig. 5.**
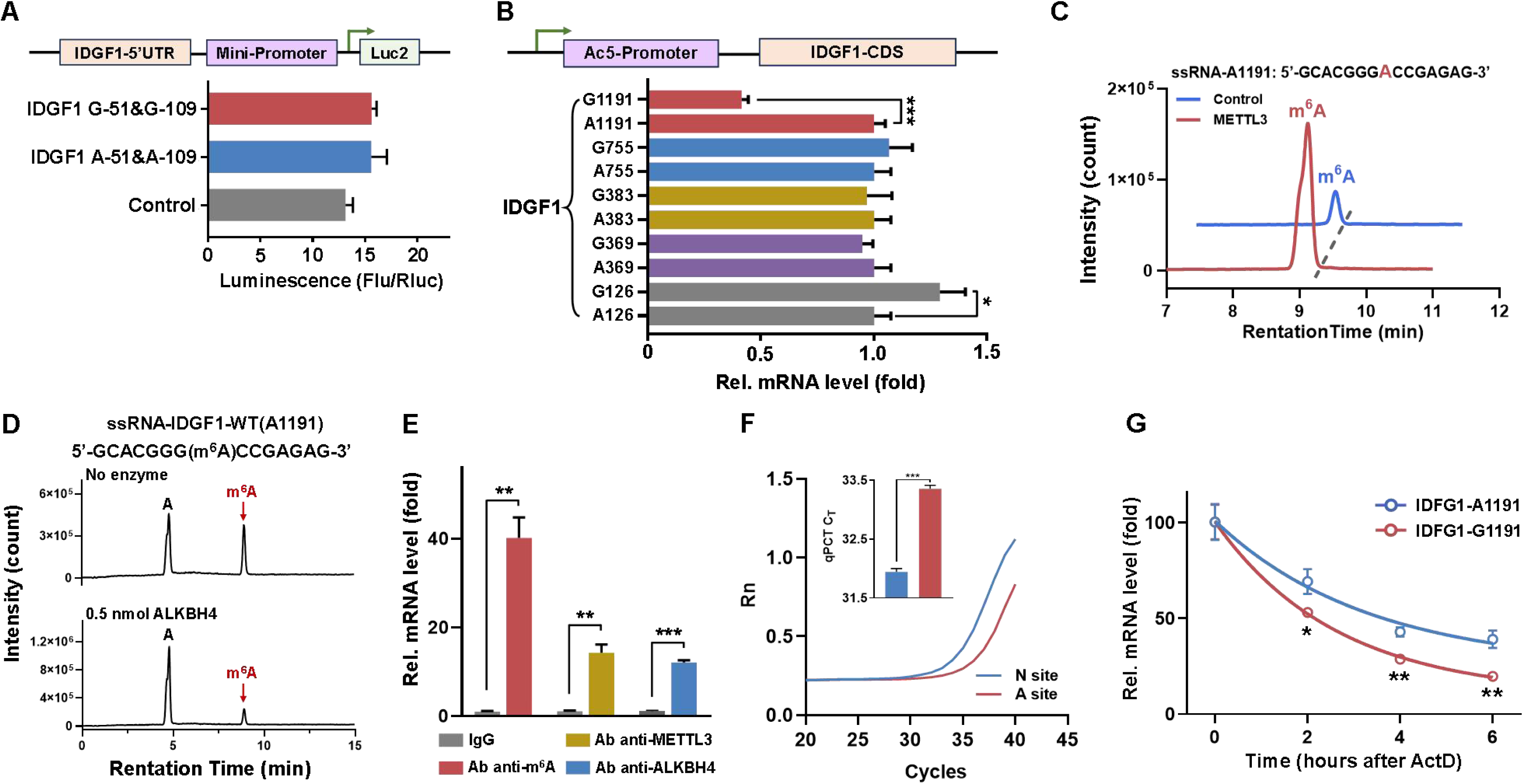
Functional characterization of the m^6^A modification site on *IDGF1* and its regulation by METTL3 and ALKBH4. (A) Identification of the activity of m^6^A binding sites in the 5’ UTR of IDGF1 constructs in dual-luciferase reporter gene assays (n = 3, means ± SE). (B) Quantification of the expression of the wild type IDGF1 CDS (IDGF1-A126, A369, A383, A755, A1191) and corresponding mutant (IDGF1-G126, G369, G383, G755, G1191) constructs by quantitative PCR following expression in *Drosophila* S2 cells. Relative mRNA levels of *IDGF1* were normalized to *Drosophila RPL32* expression (means ± SE, n = 3, \**P* < 0.05, \*\*\**P* < 0.001). (C) Activity of METTL3 (0.5 nmol) in methylating m^6^A in a ssRNA of *IDGF1* for 1 h at 25 °C as revealed by UPLC-MS/MS. (D) Activity of ALKBH4 (0.5 nmol) in demethylating m^6^A in a ssRNA of *IDGF1* at pH 7.0 and 25 °C for 30 min as revealed by UPLC-MS/MS. (E) RNA immunoprecipitation (RIP) analysis of the interaction of m^6^A antibody and METTL3 or ALKBH4 antibodies with the mRNA of *IDGF1* (n = 3, means ± SE; \*\**P* < 0.01, \*\*\**P* < 0.001, two-tailed Studentʹs *t* test). (F) Single-base resolution analysis of m^6^A sites on cenRNA in A1191 cells using the single-base elongation-and ligation-based qPCR amplification (SELECT) assay. “A site” indicates an m^6^A-modified site, whereas “N site” denotes a non-m^6^A-modified site. (means ± SE, n = 3, \*\*\**P* < 0.001). (G) mRNA stability of the wild type IDGF1 CDS (IDGF1-A1191) and corresponding mutant (IDGF1-G1191) constructs by quantitative PCR following expression in *Drosophila* S2 cells. Relative mRNA levels of *IDGF1* were normalized to *Drosophila RPL32* expression (means ± SE, n = 3, \**P* < 0.05, \*\**P* < 0.01).

To examine whether this m^6^A site can be demethylated by the m^6^A eraser, ALKBH4, we synthesized a whitefly 15-mer ssRNA (GCACGGG**m^6^A**CCGAGAG) containing the A1191 site of *IDGF1*. In vitro RNA demethylation assays determine whether specific enzymes can remove the m^6^A modification from RNA. The single-stranded oligonucleotide was incubated with an equal amount of recombinant ALKBH4 protein and m^6^A levels analyzed by UPLC-MS/MS. The m^6^A yield of the ssRNA fragment was reduced by 46% in the presence of ALKBH4 relative to the no enzyme control, further indicating that A1191 can be demethylated by ALKBH4 *in vitro* (Fig. 5D). In contrast, recombinant Bt_METTL3 enzyme was found to have the capacity to *methylate* this ssRNA *in vitro* (*P =* 0.0013, Fig. 5C, S7E). To demonstrate the role of the m^6^A site at position 1191 in the CDS of *IDGF1 in vivo*, and the action of METTL3 and ALKBH4 at this site, we assessed m^6^A abundance, and the binding of METTL3 and ALKBH4, across transcripts of *IDGF1* encompassing the A1191 site by RNA immunoprecipitation-quantitative PCR (RIP-qPCR) using antibodies for m^6^A, METTL3 and ALKBH4. RIP-qPCR is used to analyze RNA-protein interactions and study specific modifications on RNA. RNA immunoprecipitated with the m^6^A antibody was highly (40.2-fold) and significantly enriched (*P =* 0.0010, Fig. 5E) for the *IDGF1* target mRNA compared to the IgG control samples. Similarly, mRNA immunoprecipitated with the METTL3 or ALKBH4 antibodies was significantly enriched (*P =* 6.52 × 10^-4^, 14.1-fold; *P =* 3.07 × 10^-5^, 12.1-fold respectively, Fig. 5E) for the *IDGF1* target mRNA. Consistent with the RIP-qPCR results, the SELECT assay confirmed site-specific m^6^A modification on selected cenRNAs. (*P =* 5.82 × 10^-5^, Fig. 5F). Furthermore, to examine whether this m^6^A site regulates IDGF1 mRNA stability, we generated pAC5.10 constructs containing IDGF1-A1191 and its mutant IDGF1-G1191 and performed RNA decay assays. The m^6^A mutation significantly reduced the half-life of *IDGF1* mRNA (2h *P =* 1.30 × 10^-2^; 4h *P =* 1.03 × 10^-3^; 6h *P =* 1.98 × 10^-3^, Fig. 5G). These results provide clear evidence that the coding sequence of *IDGF1* contains an m^6^A site that is bound by the m^6^A writers and erasers METTL3 and ALKBH4, demonstrating that the m^6^A pathway plays a key role in the regulation of this important developmental gene (Fig. S10).

## Discussion

Our data identify ALKBH4 as an m^6^A demethylase gene in the whitefly *B. tabaci* and reveal the role of this protein, and the m^6^A pathway more generally, in the post-transcriptional regulation of key genes involved in insect development (Fig.S10). Together, these results both demonstrate that m^6^A methylation can be a dynamic and reversible process in insects and provide new insight into the functional role of this epitranscriptomic mark in arthropods.

We demonstrate that ALKBH4 can catalyze m^6^A modifications from ssRNA, dsRNA, stem-loop RNA, ssDNA and dsDNA, and its activity is iron-and α-KG-dependent. Similar metabolic activity has been reported for mammal and plant m^6^A demethylases (22, 16). While the m^6^A demethylases FTO and ALKBH5 have been identified in mammals (16, 21), in insects, previous studies have focused on m^6^A methylases and binding proteins, and the activity and function of insect demethylases remains poorly understood. Recently, however, ALKBH8 was identified as a potential m^6^A demethylase in *Aedes aegypti* and *D. melanogaster*. Incubation of recombinant ALKBH8 with total RNA extracted from Aag2 or S2 cells was correlated with a decrease in m^6^A levels in total RNA samples in dot blot assays, suggesting ALKBH8 has m^6^A demethylase activity (29). In addition, ALKBH5 of *Locusta migratoria* was identified as a putative m^6^A demethylase *in vitro*, and linked to a role in modulating aggregation behaviors (30). Finally, *TcALKBH4,* was identified in *Tribolium castaneum*, and shown to mediate the development of larvae, possibly through the 20E signaling pathway (36). However, dot blot analyses showed that RNAi knockdown of *TcALKBH4* does not seem to influence global m^6^A levels *in vivo*. Our study extends these previous works by identifying an m^6^A eraser in *B. tabaci* and demonstrating its activity *in vitro* and *in vivo*, thus confirming the reversibility of the m^6^A pathway in insects.

Numerous studied have linked m^6^A binding to the conserved DR<u>A</u>CH (D = C/A/U/G; R = A/G; H = A/C/U) consensus sequence in transcripts of mammals, plants, and insects (6, 10, 25). However, the conservation and distribution of the DRACH motif, and methylation and demethylation activity at different DRACH sites, in insects remains unresolved. Here, we demonstrated that 24 permutations of DRACH sites were less abundant overall in *B. tabaci* than in *H. sapiens*. Furthermore, while m^6^A sites were observed at highest frequency in the CDS of *B. tabaci*, as seen in *H. sapiens* and *A. thaliana,* considerably fewer sites were identified in the 3’ UTR of *B. tabaci* than in *H. sapiens*. The latter observation may explain why the epitranscriptomic regulation of m^6^A has been frequently reported to involve modification of sites occurring in the 3’ UTR region of mammalian transcripts (2, 3, 12). Regardless, our results reveal that m^6^A sites are commonly found in insect genes, prompting additional studies into the function of these sites. We show that m^6^A methylase and demethylase activity is particularly affected by the last nucleic acid of the DRACH motif. Furthermore, we reveal the conserved activity of the ALKBH proteins of *B. tabaci*, *H. sapiens*, *A. thaliana*. However, to date, a homolog of human FTO has not been identified in plants or insects (37). In relation to the mode of action of m^6^A demethylases, recent work has shown that RNA-binding motif protein 33 (RBM33) is a previously unrecognized m^6^A RNA-binding protein that activates ALKBH5 demethylase activity via suppression of ALKBH5 SUMOylation, and regulates ALKBH5-mediated m^6^A demethylation selectivity (38). Thus, further research is needed to determine whether other RBM33-like proteins influence the demethylase activity of ALKBH subfamily genes in insects and other organisms.

m^6^A methylation is the most prevalent internal RNA modification in eukaryotes, playing crucial roles in diverse biological processes (6, 20). Here, we reveal the role of m^6^A in the regulation of the chitinase-like protein IDGF1 and its impact on nymph development in *B. tabaci*. RNAi experiments revealed that *IDGF1* plays a key role in the regulation of molting at each nymphal stages, and knockdown of this gene results in increased mortality and developmental retardation. We have previously shown that 14 genes encode putative chitinase-like proteins in *B. tabaci*, of which *BtCht10*, *BtCht5,* and *BtCht7* were shown to play a key role in nymph molting (39). In another hemipteran insect species, *Sogatella furcifera*, chitinases were also shown to play crucial roles in the transition from the nymph to adult stages (40). However, epitranscriptomic regulation of chitinases has not been previously demonstrated. Rather, previous studies implicating epigenetic mechanisms in the regulation of chitinases have identified roles for DNA methylation and microRNAs. For example, Xu et al. demonstrated that DNA methylation suppressed the promoter activity of chitinase 10 via downregulation of the transcription factor homeobox protein araucan in *B. mori* to promote the development of wings (41). Additionally, the miRNA miR-282-5p was shown to regulate chitinase 5 affecting the larval moulting process in the same species (42). Here, we uncover a single m^6^A site (A1191) in the sixth exon of *IDGF1* that is bound and methylated or demethylated by the m^6^A writers and erasers METTL3 and ALKBH4 respectively. The location of the m^6^A site in *IDGF1* is consistent with previous studies which have shown that m^6^A exhibits a marked enrichment in long internal exons and near stop codons (20, 43, 44). Our findings on the role of METTL3 and ALKBH4 in *B. tabaci* development is consistent with previous studies. For example, the m^6^A methyltransferase gene *METTL3* was recently shown to play an essential role in embryonic development in silkworm, *Bombyx mori* (45). Furthermore, in *Tribolium castaneum*, m^6^A modification was shown to be required for development and reproduction (27, 36). These findings, in combination with our results on *IDGF1*, provide an emerging body of evidence of the important role of the m^6^A pathway on insect development.

In the example of m^6^A mediated gene regulation in our study m^6^A hypermethylation was associated with increased gene expression. This finding is significant as previous studies on vertebrates and plants have shown that m^6^A on mRNAs promotes their degradation in the cytoplasm leading to a post-transcriptional reduction in gene expression (1, 22). Consistent with this, work on honeybees, *Apis mellifera*, found that transcripts with high m^6^A levels tended to preferentially exhibit downregulated transcription (46). A previous study on *Arabidopsis* found that increased m^6^A methylation promoted accelerated mRNA degradation of the flowering gene *Flowering Locus T.* However, in the same study, of genes shown to contain hypermethylated peaks, 0.5% were expressed at lower levels, and 2.2% were expressed at higher levels in a line where the m^6^A demethylase ALKBH10B was knocked out compared to a wildtype line (22). This suggests that m^6^A affects RNA fate in other ways in plants that extend beyond m^6^A-dependent mRNA degradation. Furthermore, in the red flour beetle, *T*. *cataneum*, RNAi knockdown of METTL3, which would be expected to result in decreased m^6^A levels, and thus reduced mRNA degradation (and increased gene expression), resulted in the downregulation of 112 genes and the upregulation of just 11 genes. Furthermore, recent reports have revealed that m^6^A can promote cap-independent translation under conditions of stress leading to enhanced gene expression (2). Given this previous work and our findings, it is clear that further work is required to understand the roles of m^6^A on gene expression in insects and the factors influencing its action.

In summary, we demonstrate that m^6^A methylation is reversible in an insect through the action of the demethylase ALKBH4, and highlight the pivotal role of the methyltransferase-demethylase complex in regulating insect development. These findings contribute to a deeper understanding of epigenetic regulation in invertebrates.

## Supporting information

ALKBH4 supplementary information

## Acknowledgements

This research was supported by the National Natural Science Foundation of China (32221004, 32272598, 32361133558), China Agriculture Research System (CARS-24-C-02), The Beijing Key Laboratory for Pest Control and Sustainable Cultivation of Vegetables and the Science and Technology Innovation Program of the Chinese Academy of Agricultural Sciences (CAAS-ASTIP-IVFCAAS). Central Public-interest Scientific Institution Basal Research Fund (Y2023XK15; Y2024XK01).

## Author Contributions

X.Y., Y.Z., and C.B. designed the research. J.Y., C.W., P.G., X.D., B.F., S.L., X.W., C.Y., M.H., C.H., T.D., J.L., and C.Z performed the experiments and analyzed the data, X.Y., J.Y., C.B., X.G., and R.N. wrote and revised the manuscript.

