## Supplementary material for "RNA m^6^A demethylase ALKBH4 governs whitefly development": ALKBH4 supplementary information

**This PDF file includes:**

Supplementary Materials and Methods

Figures S1 to S7

Tables S1 to S8

Legends for Datasets S1 to S6

Supplemental References

**Other supplementary materials for this manuscript include the following:**

Datasets S1 to S6

**SI Materials and Methods**

**Identification of potential $m^6A$ "writers" and "erasers" in insects.** A phylogenetic tree was constructed with homologous single-copy genes of 266 insect species (genomic data taken from NCBI) by Randomized Accelerated Maximum Likelihood (RAxML) (1). Based on the reported methylases and demethylases, the prediction of conserved domains of each gene was conducted using InterPro (EMBL) and the Conserved Domain search function of NCBI. Protein family databases were used to select putative $m^6A$ methylases and demethylases by HMMER 3.1b2 (2).

**Molecular Docking.** The structures of native ligands (histidine, H; aspartic acid, D; arginine, R) and mutant ligands (alanine, A; glutamine, Q) were obtained from PubChem (NCBI). The structure of the ALKBH4 protein was predicted using AlphaFold3 (3). Docking simulations of the ligands with the protein were performed using AutoDock Vina (4), and the resulting complexes were visualized using PyMOL 2.5.

**Cloning, expression and purification of ALKBH4.** The whitefly full-length *ALKBH4* gene and two double mutants were subcloned into pET28a vector. All primers used for gene cloning are listed in Table S8. ALKBH4 and its mutants were expressed in BL21(DE3) *E. coli* and induced with 0.25 mM isopropyl β-D-1-thiogalactopyranoside (IPTG) for 12 h at 25 °C. The soluble fraction was purified by Ni-NTA column with high salt buffer at 4 °C (Beyotime™ His-tag Purification Resin). The purified proteins were analyzed by SDS-PAGE gel electrophoresis and immediately used in subsequent demethylation activity assays.

**Biochemical assay of ALKBH4 activity *in vitro*.** Demethylation activity assays were performed as previously reported (5-7). Briefly, reaction mixtures (100 μL) contained the following components: an equal amount of the recombinant protein and DNA/RNA with $m^6A$, 50 mM HEPES, 150 μM $(NH_4)_2Fe(SO_4)_2 \cdot 6H_2O$, 300 μM α-KG, 2 mM L-ascorbic acid. The reaction was incubated for 30 min at 25 °C, then quenched by heating for 5 min at 95 °C. DNA/RNA was digested with 2 μL (1 U) nuclease P1 (Sigma Aldrich) in 50 μL of buffer (20 mM NaOAc, 5 mM $ZnCl_2$, 50 mM NaCl) for 2 h at 50 °C, followed by the addition of 5 μL (1 M) of fresh $NH_4HCO_3$ and 1 μL (1 U) of alkaline phosphatase (Sigma Aldrich). The mixture was incubated for an additional 4 h at 37°C. The mixture was subsequently diluted 100-fold with 80% methanol and $m^6A$ and A nucleosides quantified by UPLC–MS/MS. UPLC–MS/MS analysis was carried out using a Waters ACQUITY UPLC I-Class/Xevo TQ-S Micro (Waters, MA, USA) equipped with a BEH C18 1.7 μm 2.1 × 50 mm column (Waters, MA, USA). Methanol was used as the mobile phase A and water as mobile phase B. The injection volume was 1 μl for each sample, and the flow rate was 0.2 mL/min. The program was set as follows: solvent A was maintained at 5% for 2 min, 15% for 8 min, 25% for 5 min, and finally 5% for 5 min. The column and FTN sample manager temperatures were set at 20°C and 4°C, respectively. The MS/MS system was set to positive ion mode for all analysis. Data analysis was performed by MassLynx V4.1 (Waters, MA, USA). The experimental UPLC–MS/MS parameters used for the determination by multiple reaction monitoring under the conditions and parameters are summarized in Table S7.

**Selecting of $m^6A$ motifs from genome.** 24 motifs were determined according to DRACH (D = C/A/U/G; R = A/G; H = A/C/U). Numbers and locations of gene IDs in each motif were determined by strictly matching the genome and transcript of each species, and non-restricted annotation information was confirmed. The code and genomic information of three species (*Bemisia tabaci*, *Homo sapiens*, *Arabidopsis thaliana*) used to screen genes are available on the site Genome websites of three species: https://github.com/YJZhang-Lab/select_genes _from _the_genome_for_each_species.

**Biochemical assay of METTL3 activity of *Homo sapiens* and *Arabidopsis thaliana and Bemisia tabaci MED* against 24 ssRNAs *in vitro*.** The full-length *METTL3* genes of *Homo sapiens* and *Arabidopsis thaliana and Bemisia tabaci MED* were cloned into pET28a vector and induced with 0.25 mM isopropyl β-D-1-thiogalactopyranoside (IPTG) for 24 h at 16 °C. Methylation activity assays were performed with slight modification as previously reported (8, 9). Briefly, reaction mixtures (100 μL) contained the following components: 50 μM of the recombinant protein, 5 μM ssRNA (24 type of ssRNAs, ssRNA of IDGF1), 5 mM SAM, 10 mM HEPES, 5 mM DTT, 50 mM NaCl, 1 mM $MgCl_2$. The reaction was incubated for 1 h at 25 °C, then quenched by heating for 5 min at 95 °C. RNA digestion to nucleosides and UPLC–MS/MS analysis are consistent with the ALKBH4 activity assays.

**Demethylation assays of *Homo sapiens* FTO, ALKBH5, *Arabidopsis thaliana* ALKBH10B and *Bemisia tabaci MED* ALKBH4 to 24 ssRNAs containing $m^6A$.** Four genes (*H. sapiens* FTO, ALKBH5, *A. thaliana* ALKBH10B and *B. tabaci* MED ALKBH4) were cloned into the pET28a vector and expression induced with 0.25 mM IPTG for 24 h at 16 °C. The expressed proteins were purified by Ni-NTA column as described above. The purified proteins were incubated with ssRNAs in 100 μL reaction mixtures for 3 h, prior to $m^6A$ detection by MS/MS as described above.

**Insect sample.** The *Bemisia tabaci* MED field strain was collected from cucumber plants in Beijing, China (40°21′ N, 116°24′ E). The strain was subsequently maintained on cotton plants under controlled conditions in a clean cage within a glasshouse. The environmental conditions were set to 25 °C, 70% relative humidity, and a photoperiod of 14 h light and 10 h dark, with no additional treatments applied.

**Transcriptome sequencing.** After feeding on a diet containing *ALKBH4* and *EGFP* dsRNA for 48 h, RNA was extracted from 100 adult whiteflies using standard TRIZOL (Invitrogen) protocols. Each group (the dsEGFP group and dsALKBH4 group) contained three biological repeats. Transcriptome sequencing was conducted on an Illumina HiSeq 4000 (Illumina, Inc., San Diego, CA, USA). Following the removal of adaptors and low-quality reads by SOAPnuke (v1.5.2) (10), the control (dsEGFP) group contained 24.9, 30.5 and 23.2 million clean reads, and the dsALKBH4 group contained 29.2, 28.3 and 25.9 million clean reads, respectively. Clean reads were located and compared with the reference genome (http://www.whiteflygenomics. org/ftp/MED/) (11) using Hierarchical Indexing for Spliced Alignment of Transcripts 2 (v2.0.4) (12). Quantitative expression analysis of each gene was performed using the StringTie software (v2.1.2) (13), and differential expression analyses were performed using the DEseq2 (v1.4.5) (14).

**Extraction of DNA and RNA and qRT-PCR in whitefly.** Genomic DNA was extracted from 70 adult whiteflies using the MiniBEST universal genomic DNA extraction kit (Takara Biotech). Total RNA was isolated from 50-60 adult whiteflies using standard TRIZOL (Invitrogen) protocols, and the quantified RNA (1 μg) converted to cDNA using oligo (dT) primer and Superscript II reverse transcriptase (Tiangen, Beijing, China). qRT-PCR was performed using 2 × RealStar Fast SYBR qPCR Mix (GenStar, Beijing, China) on the QuantStudio 5 RealTime PCR system (Applied Biosystems). Each group of whitefly adults (three repeats, n = 70) from different treatments were used for qRT-PCR analysis. Reactions comprised a total of 20 μl reaction volume containing 1 μl cDNA, 0.5 μl forward/reverse primer, and 10 μl 2 × RealStar

Fast SYBR qPCR Mix. Temperature cycling conditions were based on a two-step cycle: 2 min of activation at 95°C followed by 40 cycles of 15 s at 95°C and 30 s at 60°C. Primers with satisfactory amplification efficiencies (95%-105%) which were determined by standard curves are listed in Table S8. Two reference genes, *EF1α* and *RPL29*, were used for normalisation using the $2^{-\triangle\triangle Ct}$ method. A total of 3 biological replicates and three technical triplicates were conducted per sample.

**Nymph whitefly RNAi experiments.** Double strand RNAs (dsRNAs) were synthesized by the T7 RiboMAX Expression RNAi Kit (Promega, USA) according to the manufacturer's instructions. Enhanced green fluorescent protein (EGFP) was used to generate the control dsRNA. All primers used for producing dsRNA are listed in Table S8. RNAi on *B. tabaci* MED nymphs was performed as reported previously with slight modification (15). Briefly, the initial phase of the second nymph stage on cotton plants was used as the target for RNAi. The RNAi efficiency for *IDGF1*, *METTL3*, and *ALKBH4* is shown in Fig. S7A. The prepared 20 μL mixture contained 0.5 μg/μL *IDGF1*, *METTL3*, *ALKBN4* dsRNA, and 0.5 ng/mL nano-bioprotectant (facile-synthesized star polycation, 10 μL). The nano-bioprotectant extends the RNAi protective window, providing by Professor Jie Shen from China Agricultural University (16). Droplets containing 0.5 μg/μL dsRNA were placed on the surface of second instar *B. tabaci* MED nymphs every 48 h, and development characterised using a stereomicroscope (Leica, M205C, Germany). Mortality and developmental duration were recorded every 24 h in the second, third, and fourth instar nymphs until nymph eclosion. Each dsRNA treatment contained at least 60 independent samples of whitefly nymphs and were placed in the greenhouse with the same conditions used for rearing.

**Adult whitefly RNAi experiments.** dsRNAs were prepared as described above, and dsEGFP was regarded as the control for RNAi experiments. dsRNAs were fed to adult whiteflies by the RNAi system as previously described (17). Briefly, this RNAi system consists of feeding diet, glass tubes, Parafilm membrane, and shade cloths. The feeding diet contained 0.5 μg/μL dsRNA, 5%-volume of yeast extract and 30%-volume of sucrose. A total of 60 μL solution of feeding diet was pipetted onto the surface of a thin Parafilm membrane on the top of a glass tube, then a second layer of Parafilm was stretched over this to cover the solution. A total of 60 adult whiteflies (mixed sexes) were introduced into each glass tube for RNAi treatment, followed by collection for RNA extraction. Each treatment was performed in three biological replicates.

**VIGS assays.** Virus-induced gene silencing (VIGS) assays were conducted to investigate the effect of silencing of the *IDGF1* gene on the nymph development of *Bemisia tabaci*, and the development time and mortality rate of whiteflies from the egg stage to the adult stage were statistically recorded. A 390-bp fragment of the *IDGF1* gene was amplified from *B. tabaci* MED using gene-specific primers (Table S1) and cloned into the pTRV2 vector to generate pTRV2-*IDGF1*. Similarly, a 435-bp fragment of the *EGFP* gene was constructed into the vector (pTRV2-EGFP) as a negative control. The vectors pTRV1, pTRV2- *IDGF1*, and pTRV2-*EGFP* were introduced into *Agrobacterium tumefaciens* strain GV3101 via electroporation. Transformants were selected on LB agar plates containing 20 μg/mL rifampicin and 50 μg/mL kanamycin. PCR-confirmed *Agrobacterium tumefaciens* strains carrying *IDGF1* or *EGFP* inserts were cultured in LB medium at 28 °C with shaking at 200 rpm for expansion. Simultaneously, *A. tumefaciens* harboring the pTRV1 vector was cultured under the same conditions. When the cultures reached an $OD_{600}$ of 0.6, cells were harvested by centrifugation at 6,000 rpm for 5 min, and the supernatant was discarded. Infiltration buffer was prepared in advance by adding 40 μL of acetosyringone (AS, 200 mM), 200 μL of MES (10 mM), and 200 μL of $MgCL_2$ (10 mM) to 100 mL of sterile distilled water. The bacterial pellets were washed twice with this buffer. The resuspended cultures of pTRV1 and pTRV2-*IDGF1* or pTRV2-*EGFP* were then mixed at a 1:1 volume ratio and incubated at room temperature in the dark for 3 h. Cotton plants at the two-cotyledon stage were selected for infiltration. The abaxial side of each cotyledon was gently scratched with a 1 mL needleless syringe, taking care not to puncture through the tissue. After the dark incubation for 24 h, the plants were transferred to a growth chamber maintained at 25 °C, with a photoperiod of 16 h light/8 h dark and a relative humidity of 60%. To evaluate the effect of VIGS on the developmental performance of *Bemisia tabaci*, adult whiteflies (mixed sexes) were allowed to oviposit overnight on leaves of successfully infected VIGS cotton plants, after which all adults were removed. From the egg stage onward, nymphal development and mortality were recorded daily. A Leica microscope was used to document and compare the nymphal development of whiteflies between the treatment and control groups.

**Western blot.** Total protein was extracted from 150 adult whiteflies per sample with the cell lysis buffer for Western and IP Kit (Beyotime) following the manufacturer's instructions. Total protein and purified protein were quantified by the BCA Protein Assay Kit (Beyotime). Rabbit polyclonal antibodies were raised against a synthetic peptide of ALKBH4, IDGF1 (AtaGenix). The sequence of the peptide of ALKBH4 was H-G-D-N-S-K-Y-N-L-F-Y-S-D-H-E (from site 197 to 211); IDGF1 was E-S-R-A-S-Q-P-S-R-Q-D-L-N-E-K (from site 10 to 24). Antibody specificity was determined by BLAST search of the peptide sequence. Western blots were used to probe for *Homo sapiens* METTL3/FTO/ALKBH5 (1:10000, Abcam) and β-actin antibody (1:5000, Abcam). β-actin was used as a loading control in western blots.

**$m^6A$ Dot blot assay.** $m^6A$ dot blots were conducted as previously described with some modifications (18). Total RNA samples were denatured at 95°C for 5 minutes, and placed directly on ice. The chilled RNA was spotted on a PVDF membrane (Millipore), and dried at 90°C for 20 minutes. The membrane was blocked with 5% non-fat milk and incubated with anti-$m^6A$ antibody (1:10000, Synaptic Systems) overnight at 4°C. Then the HRP-conjugated goat anti-rabbit IgG (Beyotime) was added to the blots for 50 min at room temperature. The membrane was developed with Amersham ECL Prime Western Blotting Detection Reagent (Millipore).

**Quantitative analysis of $m^6A$ levels using UPLC-MS/MS.** UPLC-MS/MS was used to determine the $m^6A$/A ratio after RNAi knock down of whitefly methylase and ALKB family genes in *B. tabaci* MED. A total of 1 μg of total RNA extracted from *B. tabaci* adults was heated at 95 °C for 5 min, and chilled on ice immediately for 5 min. RNA was then digested by the addition of 2 μL (1 U) nuclease P1 (Sigma Aldrich) and 1 μL (1 U) of alkaline phosphatase (Sigma Aldrich). The samples were diluted 10-fold and injected into the same equipment with the same column as described above. The mobile phase A was water containing 0.1% formic acid (Fisher Chemicals), and mobile phase B was acetonitrile containing 0.1% formic acid (Fisher Chemicals). The program was set as follows: solvent A was maintained at 99% for 0.5 min, 70% for 2.5 min, 0% for 4.1 min, and finally 99% for 1.9 min.

**Cell culture and dual-luciferase reporter assays of 5'UTR of *IDGF1*.** Drosophila S2 cells were cultured in Hyclone SFX-insect medium (Thermo Scientific) at 27°C. The wild type and mutant sequence fragments of 5'UTR of *IDGF1* were cloned into the pGL4.26 reporter plasmid containing a mini-promoter. The 5'UTR of *IDGF1* was amplified from whitefly genomic DNA. All *IDGF1* fragments with m$^6$A prediction sites and mutant sites (A mutant to G) were cloned into the pGL4.26 vector. pGL4.26-IDGF1 (600 ng) and the reference reporter plasmid pGL4.73 (100 ng) were co-transfected into the S2 cells (24-well plates) using Lipofectamine 2000 (Invitrogen). Luciferase activity was detected using the Dual-Luciferase Reporter Assay System and a GloMax 96 Microplate Luminometer (Promega) after 48 h transfection (17). Luciferase activity assay was normalized to Renilla luciferase activity.

**Overexpression of coding sequences of *IDGF1* gene in *D. melanogaster* S2 Cells.** The full-length wildtype coding sequence (CDS) of *IDGF1* and mutated versions (A mutant to G) of the CDS (IDGF1-A126G; IDGF1-A369G; IDGF1-A383G; IDGF1-A755G; IDGF1-A1191G) were cloned into the pAC5.1b/V5/His expression vector (Invitrogen). The plasmids (600 ng) were transfected into S2 cells (24-well plates) using Lipofectamine 2000 (Invitrogen). qRT-PCR was used to estimate the expression of *IDGF1* after transfection into S2 cells for 48h. The relative expression levels of *IDGF1* were calculated by normalizing to a reference gene *RPL32* of *Drosophila melanogaster*. Each group was repeated three times.

**RNA Immunoprecipitation (RIP)-qPCR.** Total RNA of adult *B. tabaci* was used to conduct Methylated RNA Immunoprecipitation (MeRIP) analysis using the Magna MeRIP m$^6$A kit (17-10499, Millipore) following the manufacturer's instructions (17). The mouse IgG sample was used as a negative control to determine m$^6$A abundance of *IDGF1*, and the binding of *IDGF1* and METTL3/ ALKBH4. Briefly, 50 μg of total RNA was sheared to 100 nt in length by fragmentation buffer and purified. Anti-m$^6$A beads were subsequently incubated with m$^6$A/METTL3/ALKBH4 antibody at 25 °C for 30 min. Then purified RNA, 100 μL 5 × immunoprecipitation buffer, and 5 μL RNase inhibitor were incubated with rotation overnight at 4°C. Methylated RNA was eluted and converted to cDNA (Tiangen, Beijing, China). *IDGF1*-specific m$^6$A qPCR were used to assess m$^6$A abundance across transcripts of this gene. The primers used for these two genes RIP-qPCR are listed in Table S8. Each group was repeated three times.

**SELECT Detection.** The specific m$^6$A site on cenRNA was identified using the single-base elongation and ligation-based qPCR amplification (SELECT) assay (19, 20). Total RNA was extracted from 200 adult whiteflies using the Trizol method, and the RNA concentration was determined followed by genomic DNA removal. The SELECT reaction mixture was prepared by combining 9.8 μL of RNA template, 1.6 μL each of 0.5 μM Up and Down Probes, 2 μL of dNTPs, and 2 μL of 10× Action Buffer. The mixture was subjected to a stepwise annealing protocol: 90 °C, 80 °C, 70 °C, 60 °C, and 50 °C for 1 min each, followed by 40 °C for 6 min, and then held at 4 °C. Subsequently, the entire reaction mixture was supplemented with 0.3 μL of SELECT DNA polymerase, 0.47 μL of SELECT ligase, and 2.23 μL of ATP. The reaction was incubated at 40 °C for 20 min, followed by inactivation at 80 °C for 20 min, and then maintained at 4 °C. qPCR was subsequently performed using SELECT forward and reverse primers with the 2× RealStar Fast SYBR qPCR Mix (GenStar, Beijing, China) under the manufacturer's recommended conditions to validate the Ct values.

**mRNA stability Assays.** The full-length wildtype coding sequence (CDS) of *IDGF1* and a mutated version of the CDS (IDGF1-WT5, A1191; IDGF1-MU5, A1191G) were cloned into the pAC5.1b/V5/His expression vector (Invitrogen). The plasmids (600 ng) were transfected into S2 cells (24-well plates) using Lipofectamine 2000 (Invitrogen). After transfection for 36 h, the cells were treated with 10 mg/L actinomycin D. The cells were collected every 2 hours and extracted total RNA using TRlzol reagent. The relative expression levels of *IDGF1* were detected by qRT-PCR.

**Statistical Analysis.** The statistical significance of differences between samples was analyzed using two-tailed Student's *t*-test GraphPad 7.0. All quantitative data are reported as means ± SEM from at least three independent replicates.

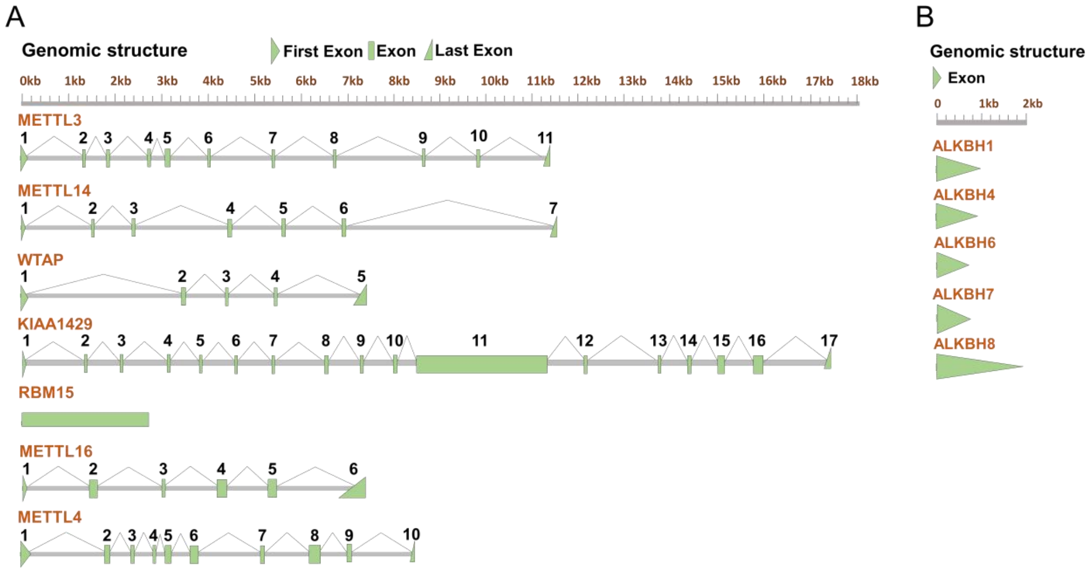


**Fig. S1.** Genomic structure of methylase and demethylase genes of *B. tabaci*. (A) The full-length *METTL3* contains a 1749 bp ORF encoding 582 amino acid residues; The full-length *METTL14* contains a 1176 bp ORF encoding 391 amino acid residues; The full-length WTAP contains a 1133 bp ORF encoding 355 amino acid residues; The full-length *KIAA1429* contains a 5703 bp ORF encoding 1900 amino acid residues; The full-length *RBM15* contains a 2274 bp ORF encoding 757 amino acid residues; The full-length *METTL16* contains a 1557 bp ORF encoding 518 amino acid residues; The full-length *METTL4* contains a 1548 bp ORF encoding 515 amino acid residues. Analysis of exon-intron structure of methylase genes indicated that *METTL3* contains 11 exons and 10 introns; *METTL14* contains 7 exons and 6 introns; WTAP contains 5 exons and 4 introns; *KIAA1429* contains 17 exons and 16 introns; *RBM15* contains 1 exon and no intron; *METTL16* contains 6 exons and 5 introns; *METTL4* contains 10 exons and 9 introns. (B) The full-length *ALKBH1* contains a 960 bp ORF encoding 319 amino acid residues; the full-length *ALKBH4* contains an 879 bp ORF encoding 292 amino acid residues; the full-length *ALKBH6* contains a 711 bp ORF encoding 236 amino acid residues; the full-length *ALKBH7* contains a 774 bp ORF encoding 257 amino acid residues; the full-length *ALKBH8* contains an 1866 bp ORF encoding 621 amino acid residues. All ALKB family genes contain a single exon.

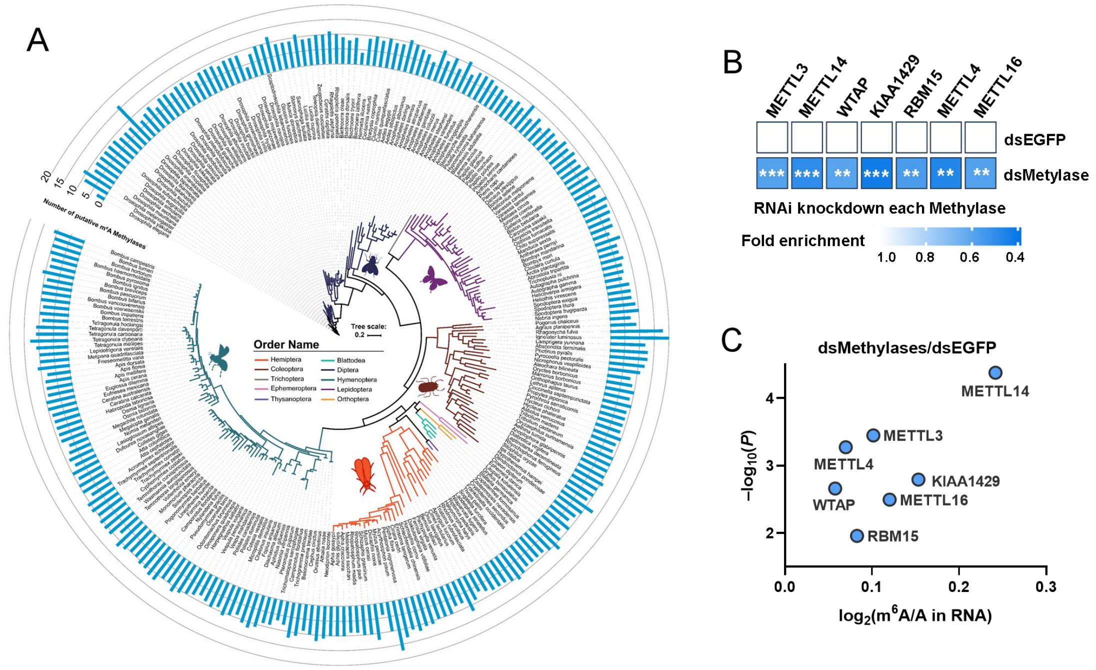


**Fig. S2. Characterization of insect and whitefly m$^6$A methylase genes.** (A) Number of putative m$^6$A methylase genes in 266 insect species. Gene numbers are presented on a maximum likelihood phylogeny of the species. Methylase genes are homologous genes of *METTL3*, *METTL14*, *METTL4* (PF05063), *METTL16* (PF05971), *WTAP* (PF17098), *KIAA1429* (PF15912), *RBM15* (PF00076). (B) Efficiency of RNAi knockdown of m6A methylase genes in *B. tabaci* MED as assessed by qRT-PCR. Relative mRNA levels were determined in *B. tabaci* adults that were fed with dsRNA at a specific concentration and time (dsMETTL3/dsMETTL14/dsMETTL4: 48 h, 0.5 μg/μL; dsWTAP: 6 h, 0.8 μg/μL; dsKIAA1429/dsRBM15/dsMETTL16: 12 h, 0.8 μg/μL). Adult whiteflies fed on dsEGFP served as the negative control (n = 3, means ± SE; ****P* < 0.001, *****P* < 0.0001, two-tailed Student's *t*-test). (C) The m$^6$A/A ratio of total RNA, determined by UPLC–MS/MS, derived from *B. tabaci* fed dsRNA corresponding to different m$^6$A methylase genes.

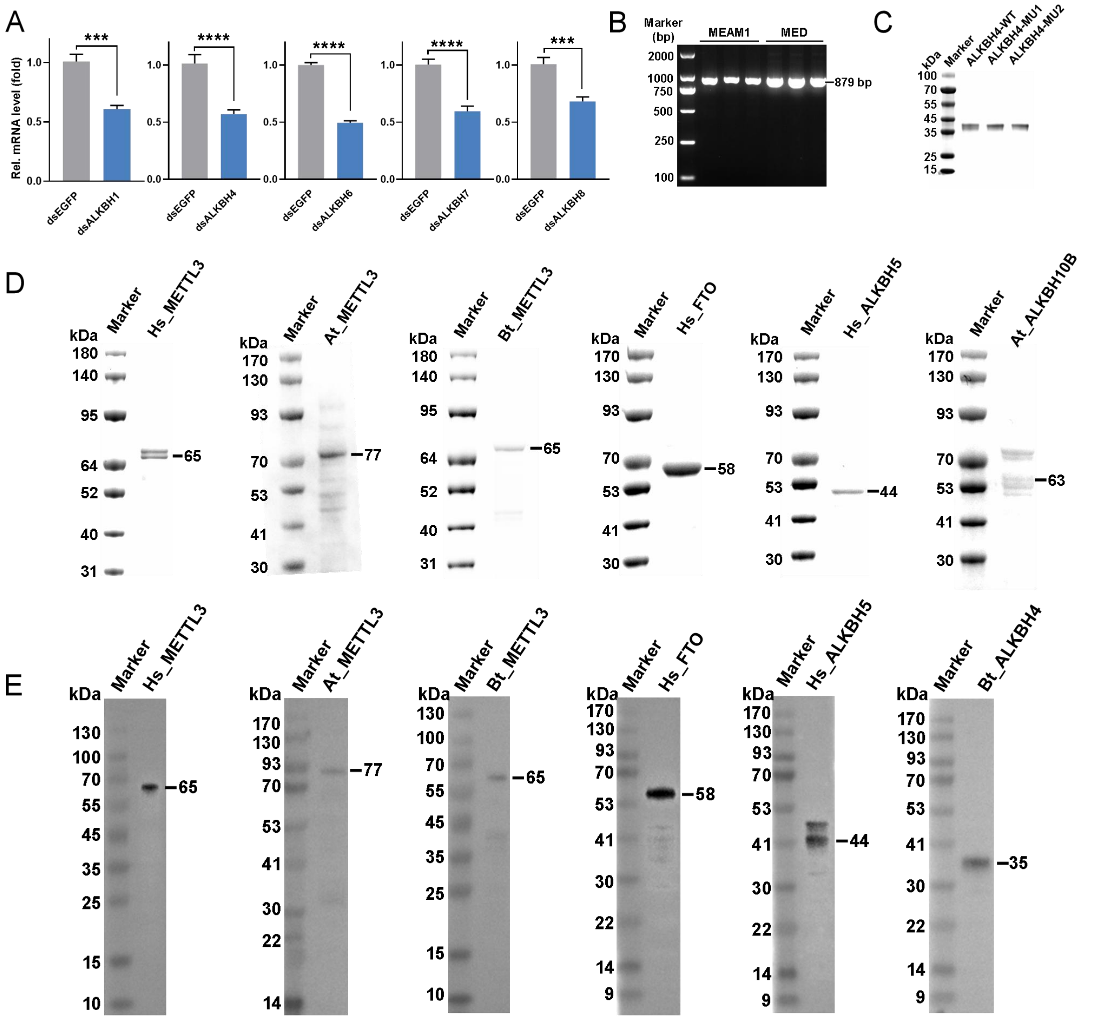


**Fig. S3.** (A) Efficiency of RNAi knockdown of ALKB family genes in *B. tabaci* MED as assessed by qRT-PCR. Relative mRNA levels were determined in *B. tabaci* adults that were fed with dsRNA at a specific concentration and time (dsALKBH1/dsALKBH8: 36 h, 0.8 μg/μL; dsALKBH4: 48 h, 0.5 μg/μL; dsALKBH6/dsALKBH7: 24 h, 0.5 μg/μL). Adult whiteflies fed on dsEGFP served as the negative control (n = 3, means ± SE; ****P* < 0.001, *****P* < 0.0001, two-tailed Student's *t*-test). (B) Full-length *ALKBH4* cloning from MEAM1 and MED whitefly. (C) Heterologous expression of the ALKBH4 wide type (ALKBH4-WT) and mutant proteins (ALKBH4-MU1/2) in *E. coli* as detected by SDS-PAGE. (D) Heterologous expression of methylase and demethylase proteins in *E. coli* as detected by SDS-PAGE. Hs_METTL3: *Homo sapiens* METTL3; At_METTL3: *Arabidopsis thaliana* METTL3; Bt_METTL3: *Bemisia tabaci* METTL3; Hs_FTO: *Homo sapiens* FTO; Hs_ALKBH5: *Homo sapiens* ALKBH5; At_ALKBH10B: *Arabidopsis thaliana* ALKBH10B. (E) Western blot analysis of heterologous expression of methylase and demethylase proteins in *E*. *coli* with specific antibodies. Hs_METTL3: *Homo sapiens* METTL3; At_METTL3: *Arabidopsis thaliana* METTL3; Bt_METTL3: *Bemisia tabaci* METTL3; Hs_FTO: *Homo sapiens* FTO; Hs_ALKBH5: *Homo sapiens* ALKBH5; Bt_ALKBH4: *Bemisia tabaci* ALKBH4.

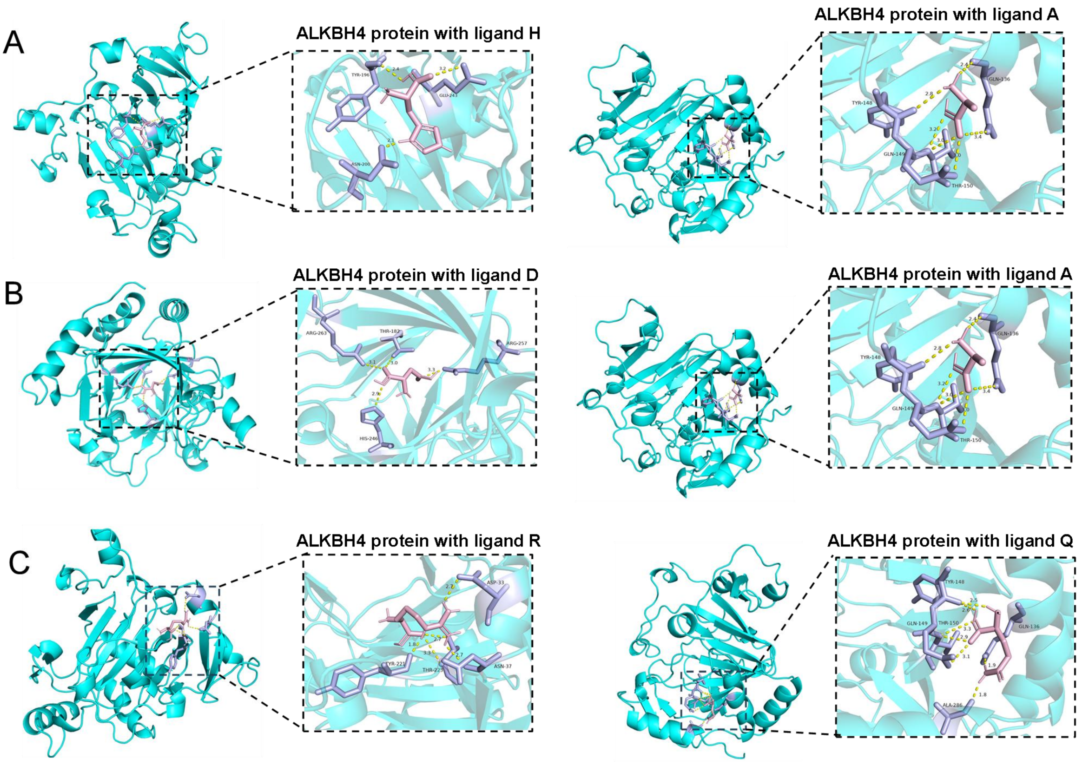


**Fig. S4. Docking of ALKBH4 protein with various ligands.** (A) The binding affinities of ALKBH4 protein with ligands H and A are –5.03 and –3.85 kcal/mol, respectively. (B) The binding affinities of ALKBH4 protein with ligands D and A are –4.61 and –3.85 kcal/mol, respectively. (C) The binding affinities of ALKBH4 protein with ligands R and Q are –5.38 and –5.0 kcal/mol, respectively.

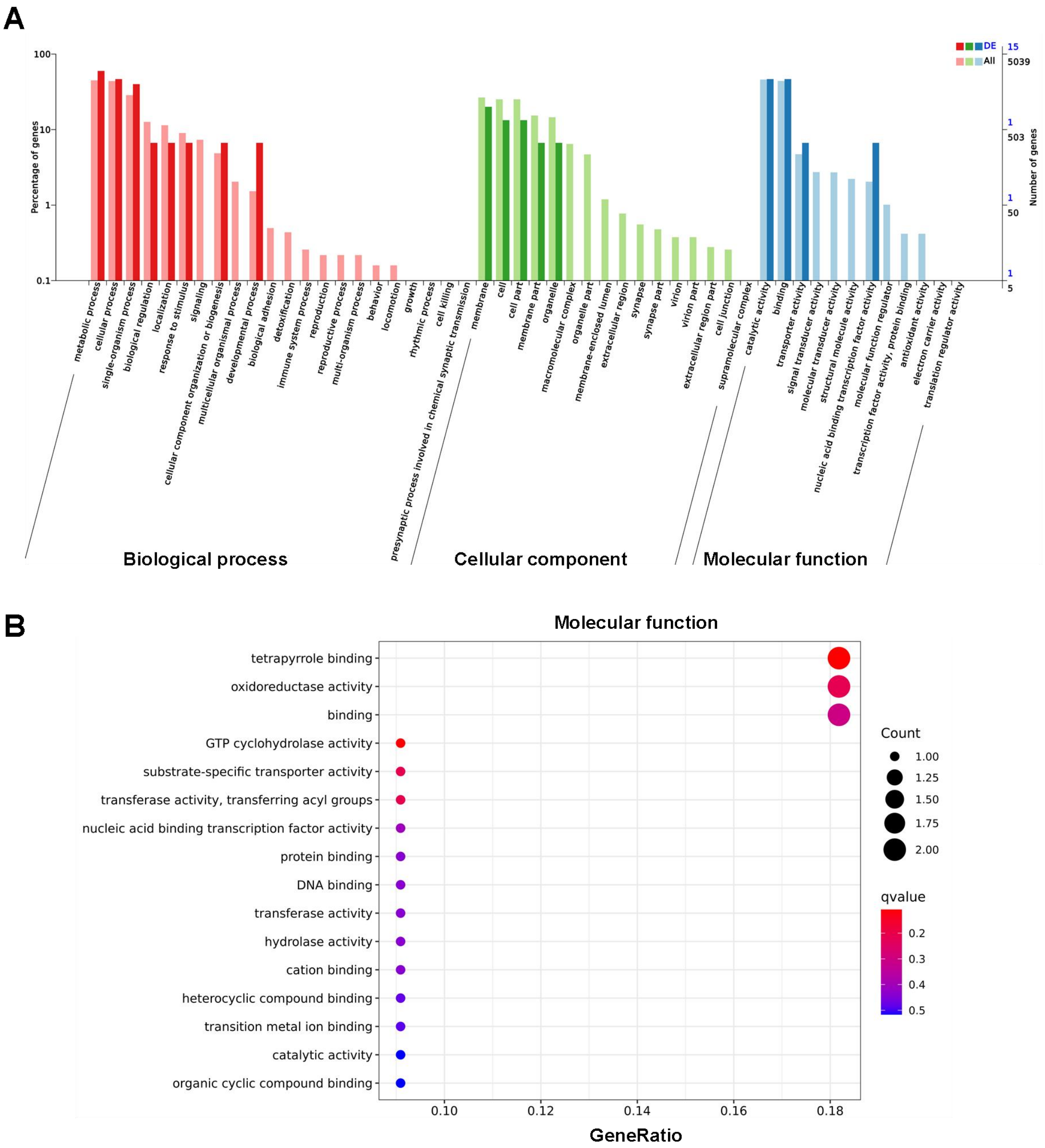


**Fig. S5.** Gene ontology (GO) enrichment analysis of differentially expressed genes identified between *B. tabaci* MED fed dsALKBH4 and dsEGFP. (A) Functional analysis of differentially expressed genes. (B) Molecular function analysis of differentially expressed genes.

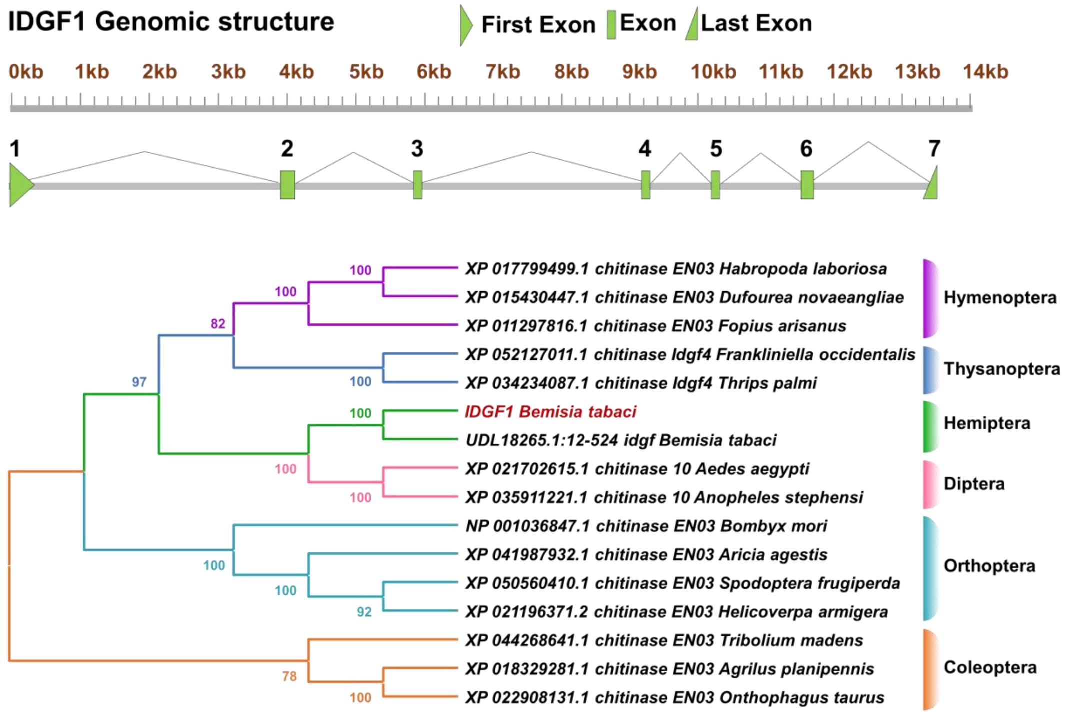


**Fig. S6.** Gene structure and phylogeny of *B. tabaci IDGF1*. The full-length cDNA sequence of *IDGF1* contained 1542-bp ORF encoding 513 amino acid residues. Genomic structure analysis indicated that *IDGF1* contains 7 exons and 6 introns.

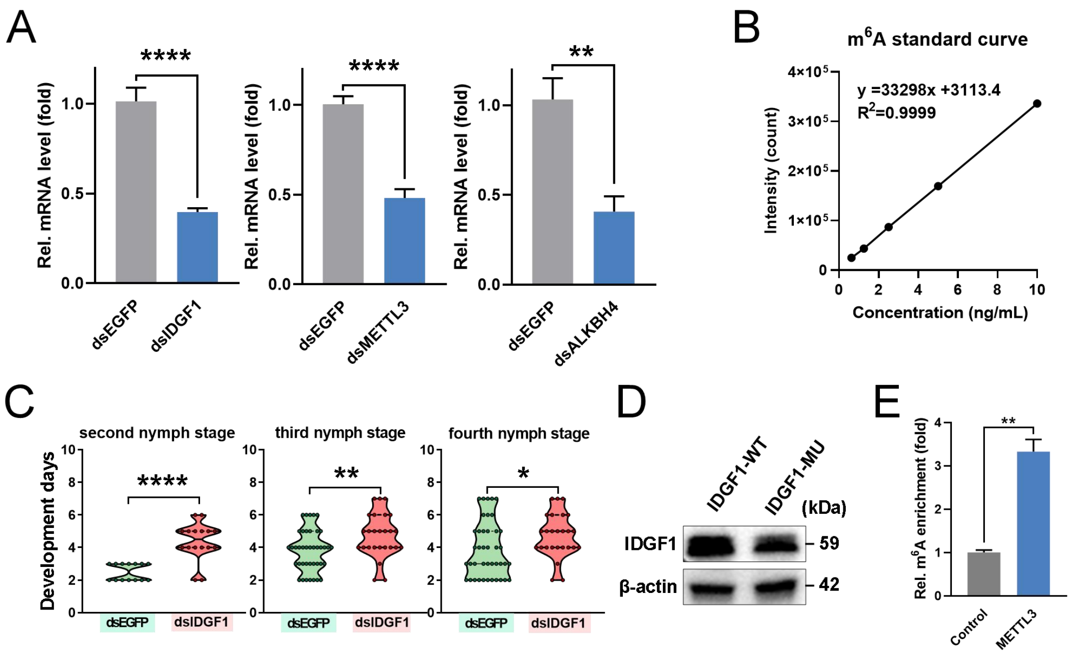


**Fig. S7.** (A) Efficiency of RNAi knockdown of *IDGF1*, *METTL3* and *ALKBH4* in the nymph stage of *B. tabaci* MED at 48 h as assessed by qRT-PCR. Relative mRNA levels were determined in the initial phase of the second nymph stage whitefly ($S^{\#1}$ strain). Nymphs exposed to dsEGFP served as the negative control (n = 3, means ± SE; $^{**}P < 0.01$, $^{****}P < 0.0001$, two-tailed Student's *t*-test). (B) An $m^6A$ standard curve was generated to quantitatively test the ALKBH4 mediated demethylation of the $m^6A$ site (A1191) in IDGF1. (C) Development of nymphs of *B. tabaci* MED following RNAi knockdown of *IDGF1* in the second, third and fourth nymphal stage; whiteflies exposed to dsEGFP served as the control ($^{*}P < 0.05$, $^{**}P < 0.01$, $^{****}P < 0.0001$, two-tailed Student's *t*-test). (D) Western blot analysis of the expression levels of the IDGF1 full-length CDS (IDGF1-WT, A1191; IDGF1-MU, A1191G) in S2 cells. β-Actin was used as a loading control. (E) Methylation was determined after the reaction of whitefly METTL3 with a 15-mer ssRNA (GCACGGGACCGAGAG), and $m^6A$ enrichment levels were measured by UPLC-MS/MS (n = 3, means ± SE; $^{**}P < 0.01$, two-tailed Student's *t*-test).

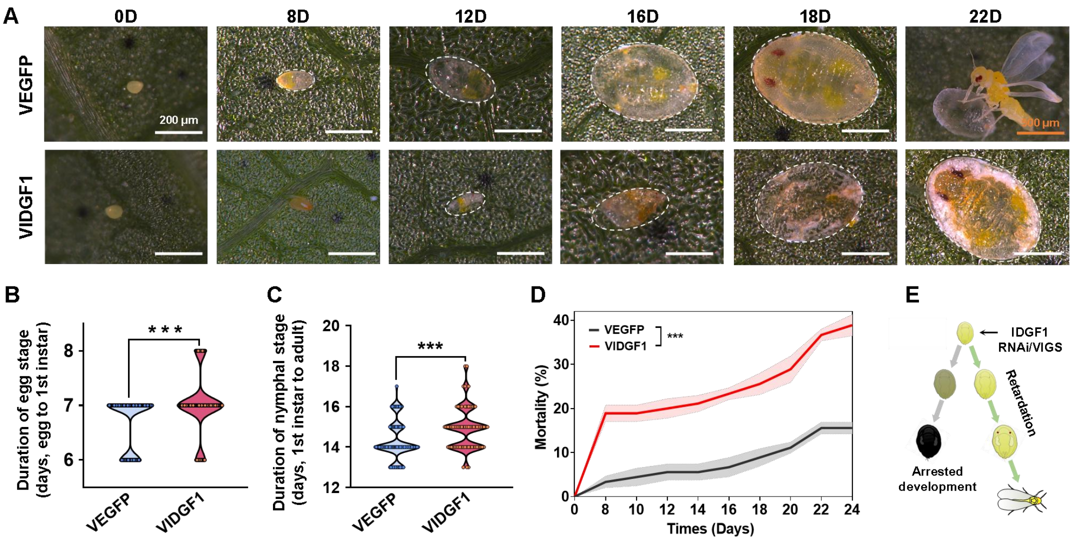


Fig. S8. (A) Representative images showing the developmental progression of *B. tabaci* following VIGS-mediated knockdown of IDGF1. Adult whiteflies were allowed to oviposit overnight on VIGS-treated cotton plants, after which all adults were removed. The offspring were monitored daily from the egg stage until death. Nymph stage: scale bars, 200 μm. Adult stage: scale bars, 500 μm. (B) Egg hatching time. (C) Nymphal developmental duration. (D) Cumulative mortality from the egg stage onward. (E) Schematic illustration of the effects of RNAi/VIGS-mediated knockdown of IDGF1 on *B. tabaci* development. Two distinct developmental phenotypes, retardation and arrested development, were observed.

CCCACTGCTCATATTAGAGCACAGAAGTAACAGAATTTTAAGTTTTTACAGTGTTTACAGCGCAAATCAATTACGCGTGTCTTCAGTCAATGTTTGAAAACTTGGCATC

5’ UTR ⟶

AATATTAAATACGTGATTCAAACCTTAGACGAGGAAAATTCAAATCGAGTCAGGCAGAATTCAAAGCCGATC**AGACT**TTCGCATCTTCAGCTGTGAATTGACATTCGA

IDGF A-51&A-109 A-109

GCGTGTTTCGATCCAAATTAAA**AGACA**TTTTCTAAAAGCGCGATGAATACCACACAGATCAAACTCGTGATTTTC**ATG**ACCATTTTTTTGAGGAACTACGTAGAGTC

A-51 CDS ⟶

GAGGGCATCACAGCCAAGCCGCCAAGATCTCAATGAAAAGTCTATCAAAGTTGAAAAACGCGAAACTCCGGGGACGAAGCACCTCCTGTCC**AGACT**GGAAAAATC

A126

AGGTATTTTCGTGCAGCCGTTACAGGAGAGGATCACCAGAAGCGAGCCGAAATTCAGGGACTTCATGGACGAGCTATCAAGCCATGGGATTTTGAAGAACAATCTA

GTGGATTCGGACCTCGCGTCGTTGACGACGGAAATCGTGAATAAAGAGATAGGAGAGGAGGACCAGCGTAAAAAGATAGTCTGCTACTATGACAGAACCAAGTCTT

ACTCAGGAAATGGTTTC**GAACT**GAATGTGAT**GGACC**TAAACGTGAGCATGAGCCTCTGCACGCACTTGGTGTACGGCTACGCGATGGTGGATCGGCAGACGGGG

A369 A383

AAGATAAGTTTGAACCACGCGGGAATGCTTCAACTCGTCGAAATCCAACACATTCAAATGATCCACGCCCTCAAGAGCTTCTATCCGAAATTCAAATTGCTTCTCGGC

ATAGGAGCGCCTCCGTCCGAGCATTATCAACACGAGAGCTACGTTCAATTCAATAACGTGCTGAGATCTGATGAGACGAGACTTCGCTTCATCAACTCGGTGGTCCC

TGTGCTGCGACGCTACAACTTCGATGGTATCGACTTGATCTGGAAGTTCGAGCGCTATCTCGAGGCCGATGATGAACACCCAAA**AGACT**GCGGAATGTGGTGTAAA

A755

TTATTGGGAGACAAGACACAGCCTGATTCGAGCAAACAAACGAATCGTTATATTGAGTTGCGGAATAGTTTTTCGGATTTGGTTCAAGAAATGAAATCAGTACTATTTA

ACTACACAATGGCCGTATCTATCCTACCCCAGATGGATGAGATATTTTACGACAAGCGAGTGATTAATTACTACGCGGATCAAATTCACCTTCTTGCATTTGACTTTATC

CTTCCATGGAACGATCTTGGAGTGGCGGAGCACACGGCACCTATCTACAAACTTGATAGTTTAGTCAATGATTGGTTGTGGCATGACGCTCAAAAGGGCAAAATCAT

AGTCGGCGTGGCAACATATGCCCGAACGTGGAGGATATCTTCCAAGAGCCTTTTCAACACCTCGCCACCAGTAATCGCCGATGGCACG**GGACC**GAGAGGCGCTTA

A1191

CAGCAAGAAAGACGGAATATTTTCTTTTTACGAGGTCTGCGTGCAGTTGGTAGAGGACGGATTCACGCAATCGATCCTCGTTCTGAACAAAGTCAACGAATCGCAAC

AATACTCGGGCACTTACGCTTATAGGTTACCGACTAATTTATCCGAGGGGCTGTGGGTAAGTTTCGATGACCCGGAATCCGCCGCCACCAAGGCAAAGTACGTCAA

GGAGAAGGGTTTAAGTGGTATGATTGTTCACGGCCTGGTTTTTGATGATTTCAAGGGAATGTGTCGAAATAGATACGACACAAGCTACCCGATCCTCCAAAGTATTGC

CAATAATTTGGTA**TGA**AAAGGCAAAACGAAAAAGCTTTTCTTACAAATGGGCCTGCTTTAGAAACTTTAAGAGTCATCTTTTTACATCAGAGGAGTAAAAATGAATTTT

3’ UTR ⟶

GTGTCCATATTTGGGTAAGAAAAATTATTCCAAGGCTATAATGTGTAGGCAGGAGAATGCGACCAGTTACTTGAGCTTGAGATAATACGTGGTTGCGGGCGGTTTTCT

AGTGGAAATAATAACGGCAGCTCGCGGTCATTTCCGGGCAACTAAAACT

**Fig. S9.** One fragment (IDGF1-A-51&A-109) including two DRACH consensus sequences in the 5’ UTR and five $m^6A$ binding sites (red) in CDS of *IDGF1* in whitefly.

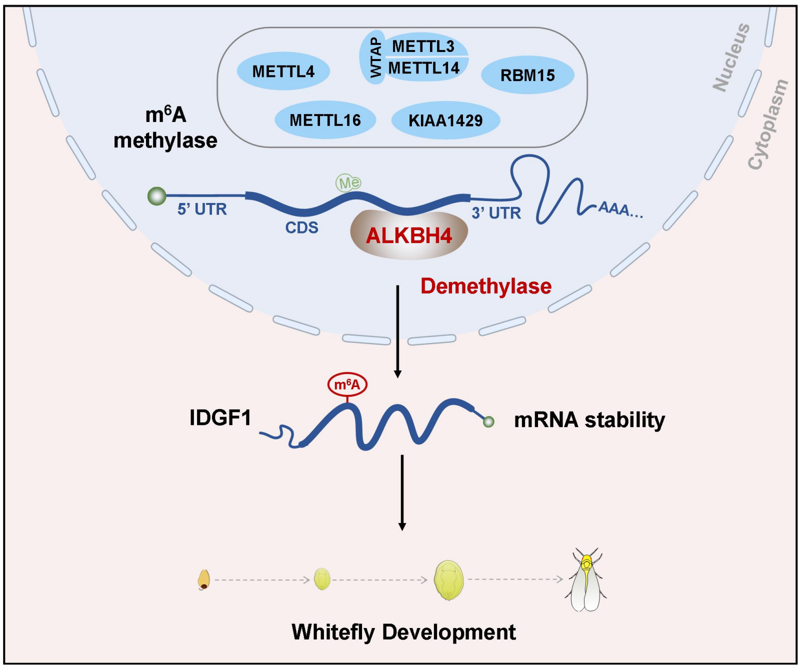


**Fig. S10**. schematic of the m$^6$A pathway in *B. tabaci*. Seven m$^6$A methylase genes and one demethylase gene are involved in the m$^6$A pathway and regulation of *IDGF1* modulating nymph development in whitefly.

**Table S1. Numbers of putative methylase and demethylase genes in the genomes of 266 insect species.**

| Name | Abbreviation | Writer | Eraser |
|---|---|---|---|
| *Aedes aegypti* | Aaegyp | 9 | 5 |
| *Anopheles albimanus* | Aalbim | 10 | 5 |
| *Aedes albopictus* | Aalbop | 11 | 5 |
| *Anopheles arabiensis* | Aarabi | 11 | 5 |
| *Anopheles atroparvus* | Aatrop | 6 | 1 |
| *Aleochara bilineata* | Abilin | 8 | 2 |
| *Anthocharis cardamines* | Acarda | 6 | 2 |
| *Atta cephalotes* | Acepha | 10 | 5 |
| *Apis cerana* | Aceran | 9 | 5 |
| *Atta colombica* | Acolom | 10 | 5 |
| *Anopheles coluzzii* | Acoluz | 10 | 5 |
| *Aphis craccivora* | Acracc | 10 | 5 |
| *Anopheles darlingi* | Adarli | 9 | 5 |
| *Apis dorsata* | Adorsa | 8 | 5 |
| *Acromyrmex echinatior* | Aechin | 8 | 4 |
| *Apis florea* | Aflore | 9 | 6 |
| *Anopheles funestus* | Afunes | 9 | 5 |
| *Anopheles gambiae* | Agambi | 8 | 2 |
| *Autographa gamma* | Agamma | 7 | 3 |
| *Aphidius gifuensis* | Agifue | 11 | 5 |
| *Anoplophora glabripennis* | Aglabr | 10 | 7 |
| *Aphis glycines* | Aglyci | 10 | 3 |
| *Aphis gossypii* | Agossy | 10 | 6 |
| *Eriosoma lanigerum* | Elanig | 11 | 3 |
| *Eufriesea mexicana* | Emexic | 10 | 5 |
| *Fopius arisanus* | Farisa | 11 | 5 |
| *Formica exsecta* | Fexsec | 11 | 4 |
| *Frankliniella occidentalis* | Foccid | 9 | 6 |
| *Frieseomelitta varia* | Fvaria | 11 | 5 |
| *Ferrisia virgata* | Fvirga | 10 | 2 |
| *Gryllus bimaculatus* | Gbimac | 10 | 4 |
| *Glossina fuscipes* | Gfusci | 12 | 5 |
| *Galleria mellonella* | Gmello | 8 | 5 |
| *Helicoverpa armigera* | Harmig | 6 | 6 |
| *Hycleus cichorii* | Hcicho | 9 | 3 |
| *Hormaphis cornu* | Hcornu | 10 | 3 |
| *Halyomorpha halys* | Hhalys | 9 | 8 |
| *Hypothenemus hampei* | Hhampe | 10 | 3 |
| *Hermetia illucens* | Hilluc | 9 | 5 |
| *Hyposmocoma kahamanoa* | Hkaham | 8 | 5 |
| *Habropoda laboriosa* | Hlabor | 9 | 5 |
| *Heliconius melpomene* | Hmelpo | 8 | 3 |
| *Hycleus phaleratus* | Hphale | 10 | 3 |
| *Harpegnathos saltator* | Hsalta | 10 | 5 |
| *Heliothis virescens* | Hvires | 8 | 3 |
| *Ignelater luminosus* | Ilumin | 20 | 3 |

| | | | | | | | |
|---|---|---|---|---|---|---|---|
| *Anopheles longipalpis* | Alongi | 9 | 3 | *Ips nitidus* | Initid | 13 | 3 |
| *Apolygus lucorum* | Alucor | 7 | 2 | *Junonia coenia* | Jcoeni | 10 | 3 |
| *Apis mellifera* | Amelli | 9 | 5 | *Lerema accius* | Lacciu | 7 | 2 |
| *Anopheles merus* | Amerus | 7 | 5 | *Lasioglossum albipes* | Lalbip | 8 | 2 |
| *Antheraea pernyi* | Aperny | 8 | 3 | *Lethrus apterus* | Lapter | 10 | 4 |
| *Acyrthosiphon pisum* | Apisum | 17 | 6 | *Lucilia cuprina* | Lcupri | 9 | 4 |
| *Agrilus planipennis* | Aplani | 9 | 5 | *Leptinotarsa decemlineata* | Ldecem | 10 | 7 |
| *Arctia plantaginis* | Aplant | 7 | 3 | *Linepithema humile* | Lhumil | 11 | 5 |
| *Autographa pulchrina* | Apulch | 7 | 3 | *Laupala kohalensis* | Lkohal | 8 | 4 |
| *Athalia rosae* | Arosae | 12 | 5 | *Lucilia sericata* | Lseric | 10 | 5 |
| *Anopheles sinensis* | Asinen | 8 | 4 | *Laodelphax striatellus* | Lstria | 11 | 2 |
| *Anopheles stephensi* | Asteph | 11 | 5 | *Lepidotrigona ventralis* | Lventr | 10 | 2 |
| *Abscondita terminalis* | Atermi | 13 | 6 | *Lamprigera yunnana* | Lyunna | 12 | 4 |
| *Amyelois transitella* | Atrans | 7 | 6 | *Marronus borbonicus* | Mborbo | 10 | 3 |
| *Abrostola tripartita* | Atripa | 7 | 3 | *Myzus cerasi* | Mceras | 12 | 3 |
| *Aethina tumida* | Atumid | 10 | 6 | *Melitaea cinxia* | Mcinxi | 8 | 6 |
| *Anopheles vaneedeni* | Avanee | 7 | 2 | *Microplitis demolitor* | Mdemol | 10 | 5 |
| *Asbolus verrucosus* | Averru | 9 | 4 | *Musca domestica* | Mdomes | 9 | 5 |
| *Blastobasis adustella* | Badust | 7 | 2 | *Megalopta genalis* | Mgenal | 8 | 6 |
| *Biston betularia* | Bbetul | 10 | 3 | *Myzus persicae* | Mpersi | 10 | 6 |
| *Bombus bifarius* | Bbifar | 10 | 4 | *Monomorium pharaonis* | Mphara | 11 | 5 |
| *Bombus breviceps* | Bbrevi | 11 | 2 | *Melipona quadrifasciata* | Mquadr | 10 | 3 |
| *Bombus campestris* | Bcampe | 11 | 2 | *Megachile rotundata* | Mrotun | 7 | 4 |
| *Bradysia coprophila* | Bcopro | 9 | 6 | *Melanaphis sacchari* | Msacch | 9 | 6 |
| *Bactrocera dorsalis* | Bdorsa | 8 | 5 | *Manduca sexta* | Msexta | 9 | 6 |
| *Blattella germanica* | Bgerma | 12 | 7 | *Nylanderia fulva* | Nfulva | 12 | 3 |

| | | | | | | | |
|---|---|---|---|---|---|---|---|
| *Bombus haemorrhoidalis* | Bhaemo | 10 | 2 | *Nebria ingens* | Ningen | 10 | 4 |
| *Bombus hortorum* | Bhorto | 10 | 2 | *Neodiprion lecontei* | Nlecon | 11 | 5 |
| *Bombus ignitus* | Bignit | 9 | 2 | *Nilaparvata lugens* | Nlugen | 10 | 4 |
| *Bombus impatiens* | Bimpat | 9 | 5 | *Nomia melanderi* | Nmelan | 9 | 4 |
| *Bactrocera latifrons* | Blatif | 4 | 5 | *Nicrophorus vespilloides* | Nvespi | 11 | 6 |
| *Bombyx mandarina* | Bmanda | 8 | 6 | *Nasonia vitripennis* | Nvitri | 9 | 5 |
| *Bombyx mori* | Bmori | 6 | 5 | *Orussus abietinus* | Oabiet | 10 | 5 |
| *Bactrocera oleae* | Boleae | 9 | 5 | *Osmia bicornis* | Obicor | 9 | 5 |
| *Bombus pascuorum* | Bpascu | 9 | 2 | *Ooceraea biroi* | Obiroi | 10 | 5 |
| *Bombus pyrosoma* | Bpyros | 9 | 5 | *Oryctes borbonicus* | Oborbo | 9 | 1 |
| *Boloria selene* | Bselen | 9 | 3 | *Odontomachus brunneus* | Obrunn | 12 | 5 |
| *Bemisia tabaci* | Btabac | 8 | 5 | *Ostrinia furnacalis* | Ofurna | 10 | 6 |
| *Bombus terrestris* | Bterre | 10 | 4 | *Orius laevigatus* | Olaevi | 12 | 2 |
| *Belonocnema treatae* | Btreat | 11 | 2 | *Osmia lignaria* | Oligna | 9 | 4 |
| *Bactrocera tryoni* | Btryon | 9 | 5 | *Oryzaephilus surinamensis* | Osurin | 10 | 3 |
| *Bombus turneri* | Bturne | 10 | 2 | *Onthophagus taurus* | Otauru | 9 | 6 |
| *Bombus vancouverensis* | Bvanco | 10 | 4 | *Pogonomyrmex barbatus* | Pbarba | 10 | 5 |
| *Bombus vosnesenskii* | Bvosne | 10 | 4 | *Polistes canadensis* | Pcanad | 10 | 5 |
| *Ceratina australensis* | Caustr | 9 | 2 | *Pogonus chalceus* | Pchalc | 7 | 3 |
| *Ceratina calcarata* | Ccalca | 9 | 5 | *Polistes dominula* | Pdomin | 9 | 5 |
| *Ceratitis capitata* | Ccapit | 10 | 5 | *Papilio glaucus* | Pglauc | 8 | 2 |
| *Cinara cedri* | Ccedri | 12 | 4 | *Pseudomyrmex gracilis* | Pgraci | 10 | 5 |
| *Cephus cinctus* | Ccinct | 9 | 5 | *Propylea japonica* | Pjapon | 9 | 3 |
| *Cyphomyrmex costatus* | Ccosta | 11 | 5 | *Papilio machaon* | Pmacha | 10 | 6 |
| *Clostera curtula* | Ccurtu | 9 | 2 | *Pieris napi* | Pnapi | 8 | 6 |
| *Cloeon dipterum* | Cdipte | 8 | 2 | *Pentalonia nigronervosa* | Pnigro | 13 | 3 |

| | | | | | | | |
|---|---|---|---|---|---|---|---|
| *Camponotus floridanus* | Cflori | 12 | 5 | *Pyrocoelia pectoralis* | Ppecto | 12 | 3 |
| *Copidosoma floridanum* | Cfloridanum | 7 | 6 | *Papilio polytes* | Ppolyt | 6 | 6 |
| *Coptotermes formosanus* | Cformo | 10 | 4 | *Pteromalus puparum* | Ppupar | 9 | 2 |
| *Colletes gigas* | Cgigas | 9 | 5 | *Photinus pyralis* | Ppyral | 13 | 7 |
| *Chelonus insularis* | Cinsul | 9 | 5 | *Pieris rapae* | Prapae | 8 | 6 |
| *Cimex lectularius* | Clectu | 9 | 6 | *Phoebis sennae* | Psenna | 8 | 3 |
| *Clunio marinus* | Cmarin | 10 | 2 | *Pyrochroa serraticornis* | Pserra | 9 | 3 |
| *Contarinia nasturtii* | Cnastu | 10 | 5 | *Pachypsylla venusta* | Pvenus | 8 | 1 |
| *Culex quinquefasciatus* | Cquinq | 9 | 5 | *Papilio xuthus* | Pxuthu | 7 | 6 |
| *Carposina sasakii* | Csasak | 9 | 2 | *Plutella xylostella* | Pxylos | 8 | 5 |
| *Cryptotermes secundus* | Csecun | 9 | 7 | *Rhynchophorus ferrugineus* | Rferru | 10 | 3 |
| *Coccinella septempunctata* | Csepte | 9 | 6 | *Rhagonycha fulva* | Rfulva | 18 | 4 |
| *Chilo suppressalis* | Csuppr | 9 | 3 | *Rhopalosiphum maidis* | Rmaidi | 9 | 6 |
| *Cotesia vestalis* | Cvesta | 10 | 3 | *Rhopalosiphum padi* | Rpadi | 10 | 3 |
| *Drosophila albomicans* | Dalbom | 9 | 5 | *Riptortus pedestris* | Rpedes | 9 | 3 |
| *Diachasma alloeum* | Dalloe | 12 | 5 | *Rhagoletis pomonella* | Rpomon | 10 | 5 |
| *Drosophila ananassae* | Danana | 16 | 5 | *Rhodnius prolixus* | Rproli | 9 | 3 |
| *Drosophila arizonae* | Darizo | 4 | 5 | *Rhagoletis zephyria* | Rzephy | 11 | 7 |
| *Drosophila biarmipes* | Dbiarm | 8 | 4 | *Sarcophaga bullata* | Sbulla | 9 | 2 |
| *Drosophila bipectinata* | Dbipec | 8 | 5 | *Stomoxys calcitrans* | Scalci | 9 | 5 |
| *Drosophila busckii* | Dbusck | 9 | 5 | *Schlechtendalia chinensis* | Schine | 11 | 3 |
| *Drosophila elegans* | Delega | 2 | 5 | *Spodoptera exigua* | Sexigu | 8 | 3 |
| *Drosophila eugracilis* | Deugra | 9 | 5 | *Sipha flava* | Sflava | 11 | 6 |
| *Drosophila ficusphila* | Dficus | 8 | 4 | *Spodoptera frugiperda* | Sfrugi | 9 | 5 |
| *Drosophila grimshawi* | Dgrims | 8 | 5 | *Sogatella furcifera* | Sfurci | 10 | 3 |
| *Drosophila guanche* | Dguanc | 9 | 5 | *Schizaphis graminum* | Sgrami | 10 | 4 |

| | | | |
|---|---|---|---|
| *Drosophila hydei* | Dhydei | 9 | 5 |
| *Drosophila innubila* | Dinnub | 9 | 5 |
| *Drosophila kikkawai* | Dkikka | 9 | 5 |
| *Drosophila mauritiana* | Dmauri | 8 | 5 |
| *Drosophila melanogaster* | Dmelan | 11 | 5 |
| *Drosophila miranda* | Dmiran | 8 | 5 |
| *Drosophila mojavensis* | Dmojav | 8 | 4 |
| *Drosophila navojoa* | Dnavoj | 10 | 4 |
| *Dufourea novaeangliae* | Dnovae | 10 | 5 |
| *Drosophila novamexicana* | Dnovam | 9 | 4 |
| *Diuraphis noxia* | Dnoxia | 4 | 4 |
| *Drosophila obscura* | Dobscu | 9 | 5 |
| *Drosophila persimilis* | Dpersi | 8 | 5 |
| *Danaus plexippus* | Dplexi | 7 | 8 |
| *Dendroctonus ponderosae* | Dponde | 9 | 5 |
| *Drosophila pseudoobscura* | Dpseud | 9 | 6 |
| *Dinoponera quadriceps* | Dquadr | 11 | 5 |
| *Drosophila rhopaloa* | Drhopa | 7 | 4 |
| *Drosophila sechellia* | Dseche | 9 | 5 |
| *Drosophila serrata* | Dserra | 7 | 5 |
| *Drosophila simulans* | Dsimul | 9 | 5 |
| *Drosophila subobscura* | Dsubob | 9 | 4 |
| *Drosophila subpulchrella* | Dsubpu | 9 | 5 |
| *Drosophila suzukii* | Dsuzuk | 9 | 4 |
| *Drosophila takahashii* | Dtakah | 8 | 5 |
| *Diabrotica virgifera* | Dvirgi | 11 | 6 |
| *Solenopsis invicta* | Sinvic | 10 | 5 |
| *Scaptodrosophila lebanonensis* | Sleban | 10 | 5 |
| *Spodoptera litura* | Slitur | 8 | 6 |
| *Sitophilus oryzae* | Soryza | 9 | 5 |
| *Stenopsyche tienmushanensis* | Stienm | 9 | 1 |
| *Tetragonula carbonaria* | Tcarbo | 11 | 1 |
| *Tribolium castaneum* | Tcasta | 10 | 6 |
| *Tetragonula clypearis* | Tclype | 13 | 2 |
| *Trachymyrmex cornetzi* | Tcorne | 11 | 5 |
| *Temnothorax curvispinosus* | Tcurvi | 11 | 5 |
| *Teleopsis dalmanni* | Tdalma | 9 | 5 |
| *Tetragonula davenporti* | Tdaven | 11 | 2 |
| *Tetragonula hockingsi* | Thocki | 9 | 4 |
| *Temnothorax longispinosus* | Tlongi | 8 | 2 |
| *Tribolium madens* | Tmaden | 9 | 6 |
| *Tetragonula mellipes* | Tmelli | 11 | 3 |
| *Trichoplusia ni* | Tni | 8 | 6 |
| *Thrips palmi* | Tpalmi | 9 | 5 |
| *Trichogramma pretiosum* | Tpreti | 12 | 5 |
| *Triatoma rubrofasciata* | Trubro | 8 | 4 |
| *Trichomalopsis sarcophagae* | Tsarco | 7 | 2 |
| *Trachymyrmex septentrionalis* | Tsepte | 11 | 2 |
| *Trachymyrmex zeteki* | Tzetek | 10 | 5 |
| *Vanessa cardui* | Vcardu | 9 | 6 |
| *Vollenhovia emeryi* | Vemery | 12 | 5 |
| *Vespa mandarinia* | Vmanda | 9 | 4 |

| | | | | | | | |
|---|---|---|---|---|---|---|---|
| *Drosophila virilis* | Dviril | 9 | 5 | *Vespula pensylvanica* | Vpensy | 8 | 4 |
| *Daktulosphaira vitifoliae* | Dvitif | 10 | 7 | *Vanessa tameamea* | Vtamea | 8 | 5 |
| *Drosophila willistoni* | Dwilli | 7 | 5 | *Vespula vulgaris* | Vvulga | 9 | 5 |
| *Drosophila yakuba* | Dyakub | 9 | 5 | *Wasmannia auropunctata* | Waurop | 11 | 5 |
| *Ephemera danica* | Edanic | 10 | 3 | *Zeugodacus cucurbitae* | Zcucur | 10 | 5 |
| *Euglossa dilemma* | Edilem | 10 | 2 | *Zootermopsis nevadensis* | Znevad | 9 | 7 |

**Table S2. Features of methylase and demethylase genes in MEAM1 and MED *B. tabaci* genomes.**

| Gene ID | Gene name | Location | Start | End | Length | Genome |
|---|---|---|---|---|---|---|
| METTL3 | OU963866.1 | Chr5 | 15211611 | 15222900 | 1749 | MEAM1 |
| METTL3 | Scaffold_79 | Chr5 | 644002 | 615222 | 1749 | MED |
| METTL14 | OU963864.1 | Chr3 | 2086617 | 2072415 | 1176 | MEAM1 |
| METTL14 | Scaffold_52 | Chr3 | 482589 | 493785 | 1176 | MED |
| WTAP | OU963862.1 | Chr1 | 45345006 | 45352464 | 1133 | MEAM1 |
| WTAP | Scaffold_209 | Chr1 | 205593 | 198137 | 1133 | MED |
| KIAA1429 | OU963865.1 | Chr4 | 52560842 | 52537560 | 5704 | MEAM1 |
| KIAA1429 | Scaffold_2874 | Chr4 | 654 | 13924 | 5704 | MED |
| RBM15 | OU963868.1 | Chr7 | 25019178 | 25021451 | 2274 | MEAM |
| RBM15 | Scaffold_222 | Chr7 | 41756 | 44029 | 2274 | MED |
| METTL4 | OU963866.1 | Chr5 | 5192202 | 5200412 | 1548 | MEAM1 |
| METTL4 | Scaffold_629 | Chr5 | 157247 | 149281 | 1548 | MED |
| METTL16 | OU963863.1 | Chr2 | 56105358 | 56112760 | 1557 | MEAM1 |
| METTL16 | Scaffold_192 | Chr2 | 303548 | 311143 | 1557 | MED |

| | | | | | | |
|---|---|---|---|---|---|---|
| ALKBH1 | OU963864.1 | Chr3 | 21857962 | 21858298 | 960 | MEAM |
| ALKBH1 | Scaffold_88 | Chr3 | 644631 | 635333 | 960 | MED |
| ALKBH4 | OU963864.1 | Chr3 | 45695902 | 45696780 | 879 | MEAM1 |
| ALKBH4 | Scaffold_282 | Chr3 | 527534 | 528412 | 879 | MED |
| ALKBH6 | OU963863.1 | Chr2 | 27512256 | 27512528 | 711 | MEAM1 |
| ALKBH6 | Scaffold_1710 | Chr2 | 113395 | 107023 | 711 | MED |
| ALKBH7 | OU963863.1 | Chr2 | 25520687 | 25521032 | 774 | MEAM1 |
| ALKBH7 | Scaffold_2681 | Chr2 | 56572 | 60472 | 774 | MED |
| ALKBH8 | OU963864.1 | Chr3 | 654686 | 655314 | 1866 | MEAM1 |
| ALKBH8 | Scaffold_1290 | Chr3 | 151102 | 141174 | 1866 | MED |

370

**Table S3. Number and proportion of m⁶A binding sites (DRACH) in the genomes of *Bemisia tabaci*, *Homo sapiens* and *Arabidopsis thaliana*.**

| Species | | *Bemisia tabaci* | | *Homo sapiens* | | *Arabidopsis thaliana* | |
|---|---|---|---|---|---|---|---|
| Type | DRACH | Numbers | Proportion (%) | Numbers | Proportion (%) | Numbers | Proportion (%) |
| 1 | CAACA | 15881 | 76.54 | 18237 | 84.80 | 22794 | 82.42 |
| 2 | CGACA | 11617 | 55.99 | 12367 | 57.50 | 14427 | 52.17 |
| 3 | AAACA | 16649 | 80.24 | 18078 | 84.06 | 24764 | 89.55 |
| 4 | AGACA | 13667 | 65.87 | 18516 | 86.09 | 21172 | 76.56 |
| 5 | UAACA | 12543 | 60.45 | 18564 | 86.32 | 20629 | 74.59 |
| 6 | UGACA | 13840 | 66.71 | 18198 | 84.61 | 19327 | 69.89 |
| 7 | GGACA | 13147 | 63.37 | 18570 | 86.34 | 16863 | 60.98 |
| 8 | GAACA | 15086 | 72.71 | 18025 | 83.81 | 21820 | 78.90 |
| 9 | CAACC | 13442 | 64.79 | 17812 | 82.82 | 17485 | 63.23 |
| 10 | CGACC | 9770 | 47.09 | 12723 | 59.16 | 10260 | 37.10 |
| 11 | AAACC | 15171 | 73.12 | 17904 | 83.25 | 23040 | 83.31 |
| 12 | AGACC | 11479 | 55.33 | 18453 | 85.80 | 15734 | 56.89 |
| 13 | UAACC | 9674 | 46.63 | 15071 | 70.07 | 16525 | 59.75 |
| 14 | UGACC | 11706 | 56.42 | 18406 | 85.58 | 14637 | 52.93 |
| 15 | GGACC | 10894 | 52.51 | 18033 | 83.85 | 12255 | 44.31 |
| 16 | GAACC | 12574 | 60.60 | 17876 | 83.12 | 18061 | 65.31 |
| 17 | CAACU | 14991 | 72.25 | 17604 | 81.85 | 20594 | 74.47 |
| 18 | CGACU | 10668 | 51.42 | 12037 | 55.97 | 12639 | 45.70 |
| 19 | AAACU | 16326 | 78.69 | 17913 | 83.29 | 24034 | 86.91 |
| 20 | AGACU | 12691 | 61.17 | 18049 | 83.92 | 19985 | 72.27 |
| 21 | UAACU | 11665 | 56.22 | 15611 | 72.59 | 19310 | 69.82 |
| 22 | UGACU | 12610 | 60.78 | 18056 | 83.95 | 19334 | 69.91 |

| | | | | | | | |
|---|---|---|---|---|---|---|---|
| 23 | GGACU | 11713 | 56.45 | 18282 | 85.00 | 15396 | 55.67 |
| 24 | GAACU | 14429 | 69.54 | 17921 | 83.33 | 19879 | 71.88 |
| Average | | **13009** | **62.70** | **17179** | **79.88** | **18373** | **66.44** |
| Total 24 motifs | | **20748** | | **21507** | | **27655** | |

**Table S4. Number of of $m^6$A binding sites (DRACH) in the 5'UTR, CDS and 3'UTR regions of genes in the genomes of *Bemisia tabaci*, *Homo sapiens* and *Arabidopsis thaliana*.**

| Species | | *Bemisia tabaci* | | | *Homo sapiens* | | | *Arabidopsis thaliana* | | |
|---|---|---|---|---|---|---|---|---|---|---|
| No. | DRACH | 5'UTR | CDS | 3'UTR | 5'UTR | CDS | 3'UTR | 5'UTR | CDS | 3'UTR |
| 1 | CAACA | 1796 | 14952 | 1735 | 18876 | 161527 | 98125 | 12944 | 92532 | 16248 |
| 2 | CGACA | 1100 | 10644 | 944 | 5877 | 41386 | 20493 | 3459 | 27191 | 4627 |
| 3 | AAACA | 2251 | 15711 | 2270 | 30335 | 187931 | 171915 | 24203 | 103037 | 28807 |
| 4 | AGACA | 1563 | 12650 | 1413 | 23734 | 165480 | 120986 | 9515 | 68238 | 11727 |
| 5 | UAACA | 1674 | 11435 | 1568 | 14473 | 86174 | 88375 | 10485 | 52649 | 13366 |
| 6 | UGACA | 1683 | 12796 | 1548 | 19951 | 147454 | 106021 | 6901 | 58330 | 9016 |
| 7 | GGACA | 1278 | 12143 | 1103 | 21703 | 152842 | 99981 | 4053 | 45096 | 4902 |
| 8 | GAACA | 1635 | 14108 | 1611 | 19038 | 143393 | 98129 | 9552 | 75208 | 12116 |
| 9 | CAACC | 1414 | 12464 | 1262 | 14154 | 112672 | 69908 | 5374 | 41924 | 6970 |
| 10 | CGACC | 925 | 8848 | 712 | 8150 | 39993 | 26358 | 2193 | 16027 | 3054 |
| 11 | AAACC | 1823 | 14208 | 1657 | 19438 | 127024 | 104317 | 15573 | 72793 | 18723 |
| 12 | AGACC | 1186 | 10463 | 986 | 20180 | 137970 | 94665 | 4364 | 34593 | 5511 |
| 13 | UAACC | 1288 | 8490 | 1132 | 9370 | 53392 | 53530 | 5962 | 31790 | 7837 |
| 14 | UGACC | 1278 | 10676 | 1153 | 19394 | 136575 | 91139 | 4020 | 30766 | 5173 |
| 15 | GGACC | 961 | 10013 | 825 | 23039 | 127850 | 88752 | 2307 | 23773 | 3140 |
| 16 | GAACC | 1278 | 11566 | 1193 | 16656 | 106043 | 74665 | 5656 | 44820 | 7612 |
| 17 | CAACU | 1724 | 14007 | 1650 | 14765 | 108883 | 77503 | 8810 | 61697 | 11021 |
| 18 | CGACU | 1084 | 9657 | 908 | 6146 | 34540 | 21615 | 3208 | 21756 | 4282 |
| 19 | AAACU | 2218 | 15423 | 2222 | 25521 | 150796 | 143454 | 18510 | 93861 | 22932 |
| 20 | AGACU | 1433 | 11720 | 1362 | 19871 | 123275 | 104210 | 10643 | 72618 | 13834 |

| | | | | | | | | | | |
|---|---|---|---|---|---|---|---|---|---|---|
| 21 | UAACU | 1650 | 10467 | 1573 | 13811 | 74284 | 87485 | 12685 | 63242 | 16570 |
| 22 | UGACU | 1557 | 11526 | 1502 | 20155 | 124730 | 106858 | 10798 | 69393 | 14277 |
| 23 | GGACU | 1155 | 10724 | 1075 | 20847 | 117068 | 93459 | 3227 | 36593 | 4346 |
| 24 | GAACU | 1679 | 13396 | 1562 | 18812 | 128514 | 98253 | 7107 | 62504 | 9325 |
| | **Total** | **35633** | **288087** | **32966** | **424296** | **2789796** | **2140196** | **201549** | **1300431** | **255416** |

**Table S5. Methylation (ug/mL) activity of METTL3 proteins from *Homo sapiens*, *Arabidopsis thaliana* and *Bemisia tabaci* against 24 ssRNAs.**

| No. | ssRNA substrate | METTL3 | | |
|---|---|---|---|---|
| | Species | *H. sapiens* | *A. thaliana* | *B. tabaci* |
| 1 | AUUGUCAACAGCAGC | 465.94±19.91 | 811.23±5.86 | 508.48±3.28 |
| 2 | AUUGUCGACAGCAGC | 452.47±21.8 | 500.34±14.36 | 364.27±16.52 |
| 3 | AUUGUAAACAGCAGC | 440.19±12.48 | 515.34±5.6 | 386.27±8.54 |
| 4 | AUUGUAGACAGCAGC | 436.13±4.36 | 481.58±0.07 | 378.93±8.66 |
| 5 | AUUGUUAACAGCAGC | 426.94±8.55 | 474.38±11.43 | 387.91±10.63 |
| 6 | AUUGUUGACAGCAGC | 395.36±15.13 | 467.2±3.26 | 404.58±11.89 |
| 7 | AUUGUGGACAGCAGC | 410.39±9.54 | 457.94±8.87 | 376.68±0.65 |
| 8 | AUUGUGAACAGCAGC | 399.36±18.96 | 464.58±8.08 | 359.89±6.52 |
| 9 | AUUGUCAACCGCAGC | 172.55±3.38 | 270.51±7.23 | 163.56±1.19 |
| 10 | AUUGUCGACCGCAGC | 169.29±2.38 | 261.96±4.88 | 158.47±0.89 |
| 11 | AUUGUAAACCGCAGC | 174.18±0.38 | 235.35±0.66 | 163.7±0.99 |
| 12 | AUUGUAGACCGCAGC | 169.28±1.13 | 256.68±1.02 | 161.57±0.3 |
| 13 | AUUGUUAACCGCAGC | 174.45±2.98 | 238.13±6.29 | 160.91±1.4 |
| 14 | AUUGUUGACCGCAGC | 170.51±3.08 | 208.14±3.88 | 162.02±1.63 |
| 15 | AUUGUGGACCGCAGC | 172.55±4.43 | 208.39±4.13 | 173.9±7.78 |
| 16 | AUUGUGAACCGCAGC | 183.58±2.83 | 237.18±0.96 | 97.46±0.5 |
| 17 | AUUGUCAACUGCAGC | 116.96±0.74 | 177.35±0.22 | 98.26±1.54 |
| 18 | AUUGUCGACUGCAGC | 111.02±1.1 | 178.22±1.56 | 96.92±1.03 |
| 19 | AUUGUAAACUGCAGC | 109.29±1.07 | 178.79±1.11 | 99.15±0.6 |
| 20 | AUUGUAGACUGCAGC | 107.73±2.39 | 188.83±0.68 | 96.96±0.93 |
| 21 | AUUGUUAACUGCAGC | 101.58±2.09 | 185.39±1.13 | 97.92±1.3 |
| 22 | AUUGUUGACUGCAGC | 98.36±0.16 | 183.08±1.02 | 101.67±1.49 |

| | | | | |
|---|---|---|---|---|
| 23 | AUUGUGGACUGCAGC | 98.66±0.28 | 191.44±0.85 | 103.83±1.47 |
| 24 | AUUGUGAACUGCAGC | 98.49±0.99 | 192.13±0.62 | 96.44±0.82 |

**Table S6. Demethylation yields (%) of demethylases from *Bemisia tabaci*, *Homo sapiens* and *Arabidopsis thaliana* against 24 ssRNAs containing m$^6$A.**

| No. | Species | *Homo sapiens* | *Homo sapiens* | *Arabidopsis thaliana* | *Bemisia tabaci* |
|---|---|---|---|---|---|
| | ssRNA substrate | FTO | ALKBH5 | ALKBH10B | ALKBH4 |
| 1 | AUUGUCA(m$^6$A)CAGCAGC | 96 ± 0.04 | 73 ± 0.05 | 62 ± 0.29 | 50 ± 0.18 |
| 2 | AUUGUCG(m$^6$A)CAGCAGC | 68 ± 0.18 | 66 ± 0.19 | 58 ± 0.23 | 16 ± 0.04 |
| 3 | AUUGUAA(m$^6$A)CAGCAGC | 92 ± 0.02 | 67 ± 0.11 | 51 ± 0.24 | 27 ±0.23 |
| 4 | AUUGUAG(m$^6$A)CAGCAGC | 96 ± 0.03 | 68 ± 0.11 | 56 ± 0.12 | 31 ± 0.34 |
| 5 | AUUGUUA(m$^6$A)CAGCAGC | 95 ± 0.01 | 67 ± 0.11 | 53 ± 0.16 | 40 ± 0.30 |
| 6 | AUUGUUG(m$^6$A)CAGCAGC | 97 ± 0.04 | 63 ± 0.31 | 53 ± 0.44 | 38 ± 0.52 |
| 7 | AUUGUGG(m$^6$A)CAGCAGC | 97 ± 0.03 | 71 ± 0.18 | 61 ± 0.22 | 39 ± 0.56 |
| 8 | AUUGUGA(m$^6$A)CAGCAGC | 95 ± 0.07 | 68 ± 0.36 | 47 ± 0.41 | 41 ± 0.68 |
| 9 | AUUGUCA(m$^6$A)CCGCAGC | 96 ± 0.02 | 11 ± 0.07 | 28 ± 0.56 | 9 ± 0.15 |
| 10 | AUUGUCG(m$^6$A)CCGCAGC | 97 ± 0.02 | 17 ± 0.34 | 34 ± 0.26 | 8 ± 0.33 |
| 11 | AUUGUAA(m$^6$A)CCGCAGC | 95 ± 0.03 | 44 ± 0.41 | 33 ± 0.37 | 21 ± 0.52 |
| 12 | AUUGUAG(m$^6$A)CCGCAGC | 97 ± 0.03 | 55 ± 0.41 | 36 ± 0.24 | 37 ± 0.24 |
| 13 | AUUGUUA(m$^6$A)CCGCAGC | 96 ± 0.01 | 24 ± 0.25 | 21 ± 0.27 | 9 ± 0.09 |
| 14 | AUUGUUG(m$^6$A)CCGCAGC | 97 ± 0.01 | 29 ± 0.35 | 39 ± 0.22 | 14 ± 0.36 |
| 15 | AUUGUGG(m$^6$A)CCGCAGC | 86 ±0.05 | 38 ± 0.23 | 26 ± 0.29 | 10 ± 0.34 |
| 16 | AUUGUGA(m$^6$A)CCGCAGC | 97 ± 0.02 | 35 ± 0.41 | 39 ± 0.23 | 11 ± 0.34 |
| 17 | AUUGUCA(m$^6$A)CUGCAGC | 98 ± 0.01 | 25 ± 0.12 | 32 ± 0.24 | 10 ± 0.28 |
| 18 | AUUGUCG(m$^6$A)CUGCAGC | 98 ± 0.03 | 31 ± 0.16 | 30 ± 0.22 | 15 ± 0.34 |
| 19 | AUUGUAA(m$^6$A)CUGCAGC | 98 ± 0.01 | 36 ± 0.28 | 42 ± 0.20 | 13 ± 0.25 |
| 20 | AUUGUAG(m$^6$A)CUGCAGC | 94 ± 0.06 | 11 ± 0.09 | 18 ± 0.22 | 10 ± 0.17 |

| | | | | | |
|---|---|---|---|---|---|
| 21 | AUUGUUA(m$^6$A)CUGCAGC | 97 ± 0.04 | 20 ± 0.54 | 38 ± 0.17 | 7 ± 0.28 |
| 22 | AUUGUUG(m$^6$A)CUGCAGC | 96 ± 0.02 | 12 ± 0.05 | 37 ± 0.25 | 20 ± 0.09 |
| 23 | AUUGUGG(m$^6$A)CUGCAGC | 96 ± 0.02 | 8 ± 0.32 | 25 ± 0.33 | 10 ± 0.28 |
| 24 | AUUGUGA(m$^6$A)CUGCAGC | 98 ± 0.04 | 27 ± 0.03 | 36 ± 0.08 | 15 ± 0.13 |

**Table S7. Experimental UPLC–MS/MS parameters used in this study.**

| Compound | Transitions (m/z) | CV (V)[a] | CE (V)[b] | DT (ms)[c] | Quantitative ion |
|---|---|---|---|---|---|
| A | 268.1+>119.1 | 40 | 40 | 0.082 | positive |
| | 268.1+>136.0 | 40 | 16 | | |
| $m^6A$ | 282.1+>123.1 | 38 | 38 | 0.082 | positive |
| | 282.1+>150.1 | 38 | 14 | | |
| dA | 252.1+>119.1 | 14 | 36 | 0.081 | positive |
| | 252.1+>136.0 | 14 | 10 | | |

CV (V)[a]: Cone voltage (V).

CE (V)[b]: Collision energy (V).

DT (ms)[c]: Dwell time (ms).

**Table S8. Oligonucleotide primers used in this study.**

| Genome ID (MEAM1) | Primer name | Primer sequence (5′-3′) | Annealing temperature (°C) | Product length (bp) | Purpose |
|---|---|---|---|---|---|
| Bta14570 | METTL3-F | ATGTCAGATGCATGGGAGGATATC | 60 | 1749 | Gene clone |
| | METTL3-R | TCATGATTTGGGAGGCACCAT | 59 | | |
| | METTL3-F | TCTGCGGACACTTTAGGCATTAT | 63 | 177 | qRT-PCR (99%) |
| | METTL3-R | AAATAACGATGTGACTGGCAATG | 61 | | |
| | METTL3-F | TAATACGACTCACTATAGGGAACAGGACGGACTGGTCAT | 57 | 326 | RNAi |
| | METTL3-R | TAATACGACTCACTATAGGCACCATGCAATTTCCATCAG | 57 | | |
| Bta06806 | METTL14-F | ATGAGTATGTTGCGAGAGTTGAAAG | 65 | 1176 | Gene clone |
| | METTL14-R | TTAACGTCCTCTGCCTCTACCTC | 63 | | |
| | METTL14-F | ATAGGTGAAGTAGCAGCCGCAAG | 62 | 141 | qRT-PCR (101%) |
| | METTL14-R | GGTTGTGTTTGTTCGTATCCAGC | 63 | | |
| | METTL14-F | TAATACGACTCACTATAGGAACGGACAAAGGAACATTGC | 57 | 332 | RNAi |
| | METTL14-R | TAATACGACTCACTATAGGTTCGATACGTTCAGTGCAGC | 57 | | |
| Bta14734 | WTAP-F | GCGTACGTTTTCGTTTTTTAAGGTT | 63 | 1133 | Gene clone |
| | WTAP-R | GATCAATCCCTGGTTGCGGTT | 63 | | |
| Bta14879 | KIAA1429-F | GTGTTTACAGTGATTTTCTTTGGCC | 62 | 5850 | Gene clone |
| | KIAA1429-R | ACCTGGGGAATGGACGTAAGAG | 62 | | |
| Bta10676 | RBM15-F | ATGAATGAAGTGTCGGGCAAGT | 61 | 2535 | Gene clone |
| | RBM15-R | TGAGATGATTCAAGCCGTGCC | 63 | | |
| Bta01886 | METTL4-F | GTGCGCGCGCTCTTACTGA | 63 | 1532 | Gene clone |
| | METTL4-R | CGTTTCGTCCATTGCCCGT | 63.7 | | |
| Bta13474 | METTL16-F | ATGCATCCCAGAAATATCTATCGTA | 51 | 1554 | Gene clone |
| | METTL16-R | GATTGTATTTTGTTTCATATTATTTC | 53 | | |
| Bta06166 | ALKBH1-F | ATGACTTTTCGGGATAAATTCAAAT | 60 | 972 | Gene clone |
| | ALKBH1-R | TCAAGCCTCCTTTAAGAGAGTTTCC | 62 | | |
| | ALKBH1-F | GATTTGAGGAGGTTCTGGATTTTTC | 62 | 185 | qRT-PCR (100%) |
| | ALKBH1-R | TGTGAAAGGGTTGCGGATAAATA | 62 | | |
| | | TAATACGACTCACTATAGGGACTCAGCCGAGTGGTCATT | 57 | 328 | RNAi |

| | | | | | |
|---|---|---|---|---|---|
| | ALKBH1-F | | | | |
| | ALKBH1-R | TAATACGACTCACTATAGGGAAGTTCGATTTTCGTCCCA | 57 | | |
| | ALKBH4-F | ATGGACCGATCCAACTTTTGTG | 61 | 879 | Gene clone |
| | ALKBH4-R | TCAAGTGAGGAAAAAGTTCTTTGCT | 61 | | |
| Bta07796 | ALKBH4-F | GTGATTCGGTTCTTACGCTTCG | 62 | 175 | qRT-PCR (102%) |
| | ALKBH4-R | TCCCATTCATACCTGGCCTCA | 63 | | |
| | ALKBH4-F | TAATACGACTCACTATAGGTAAAGGCGTCAGGACTTGCT | 57 | 456 | RNAi |
| | ALKBH4-R | TAATACGACTCACTATAGGGCCGGGTCATATTCAAGAGA | 57 | | |
| | ALKBH6-F | GCTCATCCTAGGATTGTCACATCTC | 61 | 818 | Gene clone |
| | ALKBH6-R | ATGCCAATTGGTCATTTTATCACAT | 62 | | |
| Bta10915 | ALKBH6-F | AAGAAGATATGTACCATCGCCACCT | 63 | 173 | qRT-PCR (99%) |
| | ALKBH6-R | CTTTATTTTGGTACTCTTTGGTGCG | 62 | | |
| | ALKBH6-F | TAATACGACTCACTATAGGACCGAGGTTCCTCCAAAAGT | 57 | 348 | RNAi |
| | ALKBH6-R | TAATACGACTCACTATAGGAGATCCGCAGGTTATGGTCG | 57 | | |
| | ALKBH7-F | GCCATATGATGCTTTTTAGTCACTG | 61 | 789 | Gene clone |
| | ALKBH7-R | GACTGAAGACTCAGAGGTTAGACGG | 61 | | |
| Bta10992 | ALKBH7-F | ATGAGGAATGCGATGCGTTAT | 60 | 94 | qRT-PCR (99%) |
| | ALKBH7-R | TTCTTCTTGATTTGTGGACTTGTTC | 60 | | |
| | ALKBH7-F | TAATACGACTCACTATAGGATGGTGAAACGCCTCACTTC | 57 | 409 | RNAi |
| | ALKBH7-R | TAATACGACTCACTATAGGTATTGCCGCAAAATCGTACA | 57 | | |
| | ALKBH8-F | ATGGAAGATCTACCTCTTTTGTATCTTG | 61 | 1863 | Gene clone |
| | ALKBH8-R | GCTCTTCACTAAGATGACACACCAA | 61 | | |
| Bta06335 | ALKBH8-F | AAGTCGTCCTGAACTGTCCTTATGG | 63 | 138 | qRT-PCR (99%) |
| | ALKBH8-R | GAAGCATTCAACGAAAGAGACAAGA | 62 | | |
| | ALKBH8-F | TAATACGACTCACTATAGGTATCCATGTCACCAGGCAAA | 57 | 372 | RNAi |
| | ALKBH8-R | TAATACGACTCACTATAGGGCGGGATACTTTCCATCAGA | 57 | | |
| | IDGF1-F | TCAAACTCGTGATTTTCATGACCAT | 63 | 1618 | Gene clone |
| | IDGF1-R | ACTCCTAAAGTTTCTAAAGTTGCTCCA | 61 | | |
| | IDGF1-F | CCGTCCGAGCATTATCAGCA | 63 | 134 | qRT-PCR (99%) |
| | IDGF1-R | ACCGAGTTGATGAAGCGAAGTCT | 63 | | |
| Bta02273 | IDGF1-F | TAATACGACTCACTATAGGATGGAACGATCTTGGAGTGG | 57 | 390 | RNAi |
| | IDGF1-R | TAATACGACTCACTATAGGTCCGGGTCATCGAAACTTAC | 57 | | |
| | IDGF1-F | ACGCTCAAAAGGGCAAAATCATA | 63 | 169 | RIP-qPCR |
| | IDGF1-R | ACGCAGACCTCGTAAAAAGAAAATA | 61 | | |
| Bta04011 | EF1α-F | TAGCCTTGTGCCAATTTCCG | 60 | 110 | qRT-PCR (103%) |
| | EF1α-R | CCTTCAGCATTACCGTCC | 60 | | |
| Bta02466 | RPL29-F | TCGGAAAATTACCGTGAG | 60 | 144 | qRT-PCR (101%) |
| | RPL29-R | GAACTTGTGATCTACTCCTCTCGTG | 60 | | |
| | Dm RPL32-F | GACAGTATCTGATGCCCAACATC | 58.7 | 170 | qRT-PCR (100%) |
| | Dm RPL32-R | CTTCTTGGAGGAGACGCCGT | 61.6 | | |

F: forward primer; R: reverse primer.

Whitefly Genome Database: http://www.whiteflygenomics.org/cgi-bin/bta/index.cgi

**Other Supplementary Material for this manuscript includes the following:**

**Dataset S1.** Sequences of putative methylases of 266 insects.

**Dataset S2.** Sequences of putative demethylases of 266 insects.

**Dataset S3.** Gene IDs, location annotations and NR annotations were screened from the genome of Whitefly *B. tabaci*.

**Dataset S4.** Gene IDs, location annotations and NR annotations were screened from the genome of *Arabidopsis thaliana*.

**Dataset S5.** Gene IDs, location annotations and NR annotations were screened from the genome of *Homo sapiens*.

**Dataset S6.** The differentially expressed genes derived from transcriptome sequencing after RNAi knock down of ALKBH4 in *B. tabaci* MED.
